# A computational approach to understanding effort-based decision-making in depression

**DOI:** 10.1101/2024.06.17.599286

**Authors:** Vincent Valton, Anahit Mkrtchian, Madeleine Moses-Payne, Alan Gray, Karel Kieslich, Samantha VanUrk, Veronika Samborska, Don Chamith Halahakoon, Sanjay G. Manohar, Peter Dayan, Masud Husain, Jonathan P. Roiser

**Affiliations:** Institute of Cognitive Neuroscience, University College London, London, UK; Division of Psychiatry and Max Planck Centre for Computational Psychiatry and Ageing Research, Queen Square Institute of Neurology, University College London, London, UK; Department of Clinical, Educational and Health Psychology, University College London, London, UK; Nuffield Department of Clinical Neurosciences and Department of Experimental Psychology, Oxford University, Oxford, UK; Max Planck Institute for Biological Cybernetics and the University of Tübingen, Tübingen, Germany

**Keywords:** Effort-based decision-making, Depression, Computational Psychiatry, Motivation, Anhedonia

## Abstract

**Background:** Motivational dysfunction is a core feature of depression, and can have debilitating effects on everyday function. However, it is unclear which disrupted cognitive processes underlie impaired motivation, and whether impairments persist following remission. Decision-making concerning exerting effort to obtain rewards offers a promising framework for understanding motivation, especially when examined with computational tools which can offer precise quantification of latent processes.

**Methods:** Effort-based decision-making was assessed using the Apple Gathering Task, where participants decide whether to exert effort via a grip-force device to obtain varying levels of reward; effort levels were individually calibrated and varied parametrically. We present a comprehensive computational analysis of decision-making, initially validating our model in healthy volunteers (N=67), before applying it in a case-control study including current (N=41) and remitted (N=46) unmedicated depressed individuals, and healthy volunteers with (N=36) and without (N=57) a family history of depression.

**Results:** Four fundamental computational mechanisms that drive patterns of effort-based decisions, which replicated across samples, were identified: overall bias to accept effort challenges; reward sensitivity; and linear and quadratic effort sensitivity. Traditional model-agnostic analyses showed that both depressed groups showed lower willingness to exert effort. In contrast with previous findings, computational analysis revealed that this difference was primarily driven by lower effort acceptance bias, but not altered effort or reward sensitivity.

**Conclusions:** This work provides insight into the computational mechanisms underlying motivational dysfunction in depression. Lower willingness to exert effort could represent a trait-like factor contributing to symptoms, and might represent a fruitful target for treatment and prevention.

## Introduction

Motivational impairment is common in depression (Fervaha, Foussias, Takeuchi, Agid, & Remington, 2016; Pelizza & Ferrari, 2009; Yuen et al., 2015), closely linked to anhedonia, decision-making difficulties and fatigue, which together predict poorer treatment outcome (Uher et al., 2012) and lower quality of life (Spijker, Bijl, De Graaf, & Nolen, 2001). Although anhedonia—the loss of interest or pleasure in previously enjoyable activities—is a cardinal symptom of depression, in-the-moment pleasurable experience appears to be relatively preserved in depressed individuals (Amsterdam, Settle, Doty, Abelman, & Winokur, 1987; Beevers, Strong, Meyer, Pilkonis, & Miller, 2007; Bylsma, Taylor-Clift, & Rottenberg, 2011; Chentsova-Dutton & Hanley, 2010; Dichter, Smoski, Kampov-Polevoy, Gallop, & Garbutt, 2010; Peeters, Nicolson, Berkhof, Delespaul, & deVries, 2003); by contrast, reward learning and decision-making have repeatedly been shown to be impaired in depression (Halahakoon et al., 2020), consistent with disrupted motivation.

A consistent theme emerging from this body of work is that *effort-based decision-making for reward* offers a promising lens through which motivational dysfunction can be understood. A ubiquitous finding is that the willingness to engage in effort (physical or mental) depends on both the perceived reward magnitude, and the discounting of that reward by the effort required to obtain it (Chong et al., 2017). While numerous studies have examined effort discounting in healthy individuals (Bonnelle, Manohar, Behrens, & Husain, 2016; Chong et al., 2017; Croxson, Walton, O’Reilly, Behrens, & Rushworth, 2009; Klein-Flügge, Kennerley, Friston, & Bestmann, 2016; Kurniawan, Guitart-Masip, Dayan, & Dolan, 2013; Pessiglione, Vinckier, Bouret, Daunizeau, & Le Bouc, 2018), and contemporary neurocognitive models of depression suggest that lower willingness to exert effort drives depressive symptoms related to motivation (Eshel & Roiser, 2010; Pizzagalli, 2022; Roiser, Elliott, & Sahakian, 2012), few studies have investigated these processes in patients experiencing motivational symptoms.

The first systematic attempt to examine effort-based decision-making in depression (i.e., measuring *explicit choices* to exert effort, as opposed to the degree of exertion (Cléry-Melin et al., 2011)) was reported by Treadway and colleagues (2012), who compared performance on the *Effort Expenditure for Rewards Task* (EEfRT; (Treadway, Buckholtz, Schwartzman, Lambert, & Zald, 2009) between currently depressed individuals and healthy volunteers. Drawing on an extensive literature in rodents (Salamone, Yohn, López-Cruz, San Miguel, & Correa, 2016), the EEfRT requires participants to choose between a “hard” versus an “easy” task (quick versus slow button pressing), with more reward delivered for hard choices, with varying levels of probabilities. Treadway and colleagues reported that depressed participants were less willing to choose the hard task, and less sensitive to both reward and probability. Since this initial study, several others have reported comparable results in currently depressed individuals (Yang et al., 2016; Yang et al., 2014; Zou et al., 2020), although with discrepant results for remitted depression (Berwian et al., 2020; Yang et al., 2014).

Although these studies provided an important foundation for advancing our understanding of disrupted motivation in depression, several design features complicate their interpretation. First, most studies, including those examining remitted depression, recruited individuals taking antidepressant medication, which represents a potentially important confound given that antidepressants are known to blunt neural reward processing (McCabe, Mishor, Cowen, & Harmer, 2010). Second, the exertion level required to obtain reward was typically not calibrated individually; making the hard task easier for some participants than others. Third, the inclusion of different probability conditions (with no deterministic condition) raises the possibility of an interaction between probability and effort discounting. Fourth, effort levels are not varied parametrically in the EEfRT, and no high-reward/low-effort or low-reward/high-effort options are included. Consequently, individual differences in choices of the high-reward/high-effort option could be driven by sensitivity to either reward or effort.

Another effort-based decision task, the Apple Gathering Task (AGT(Bonnelle et al., 2015), offers several key advantages: a grip-strength device is used for exertion (Cléry-Melin et al., 2011; Schmidt et al., 2008), with force level calibrated to each participant; reward and effort are varied independently and parametrically; and outcomes are deterministic. Studies using the AGT have reported that patients with Parkinson’s disease (PD; in which depression is very common (Costello, Husain, & Roiser, 2024)) were less willing to exert effort for low rewards (Chong et al., 2015), especially those with pronounced apathy (Le Heron et al., 2018). Other studies found that experimentally induced inflammation increased sensitivity to effort in healthy volunteers (Draper et al., 2018), and that, relative to matched controls, patients with treatment-resistant schizophrenia were both less willing to exert effort overall and less sensitive to reward (Saleh et al., 2023). However, no study to date has reported behaviour on the AGT in depression.

Over the past decade, computational modelling has provided important insights into the cognitive processes underlying disrupted motivation (Adams, Huys, & Roiser, 2016). A computational approach systematically evaluates competing models to identify the latent cognitive processes governing behaviour. Computational modelling enables the precise measurement of cognitive processes by estimating specific parameters, offering deeper insights than traditional (model-agnostic) measures of performance, often with superior psychometric properties (Karvelis, Paulus, & Diaconescu, 2023; Mkrtchian, Valton, & Roiser, 2023).

Therefore, in the present study we performed a computational analysis of AGT performance to provide insights into the cognitive processes underlying effort-based decision-making in depression. To address the question of whether altered effort-based decision-making is merely a consequence of ongoing depressive symptoms, or if it plays a causal role in their development, we studied four groups, all unmedicated: currently depressed individuals (MDD); individuals remitted from depression (REM); healthy individuals with a depressed close relative, but no personal history (REL, who are known to be at elevated risk); and healthy individuals without any personal or family history of psychiatric diagnosis (CTR).

Drawing on contemporary neurocognitive models of depression (Eshel & Roiser, 2010; Pizzagalli, 2022; Pringle, McCabe, Cowen, & Harmer, 2013; Roiser et al., 2012), we hypothesised that lower willingness to exert effort drives depressive symptoms, especially those related to motivation. We predicted that the MDD group would accept fewer offers to exert effort relative to the CTR group, and that this would be driven by lower sensitivity to reward and greater sensitivity to effort. Including the REL and REM groups allowed us to examine whether effort-based decision-making represents a risk factor for depression. If indeed it is a trait marker and risk factor, any motivational impairment observed in currently depressed individuals should also be present in the REM group, and possibly also the REL group (albeit they are at lower risk of developing a future depressive episode than the REM group). Given this, we predicted that a similar pattern of behaviour would be observed in the REM group as in the MDD group, and also conducted comparisons including the REL group.

## Methods

### Participants

Two studies were conducted, an initial pilot with healthy volunteers (HVs; “Pilot”), and a study including four groups (“Case-control”). All participants were aged 18-60 years, unmedicated and native English speakers. They were recruited through local advertisements, institutional participant databases and local outpatient psychological treatment services, and all provided written informed consent. The study received ethical approval from the UCL Research Ethics Committee (fMRI/2013/005) and the London Queen Square NHS Research Ethics Committee (for depressed individuals: 10/H0716/2).

#### Pilot study

Only HVs (N=102) completed this study. The final sample consisted of 67 participants (see Supplemental Information [SI] for exclusions).

#### Case-control study

Sixty-two HVs without a family history of depression (CTR; independent from the Pilot study), 38 HVs with a depressed first-degree relative (REL), 50 remitted depressed participants (REM), and 51 currently depressed participants (MDD) completed the Case-control study. The final sample included 57 CTR, 36 REL, 46 REM and 41 MDD participants (see SI for exclusions). With this sample size, at alpha=0.05, we had 80% power to detect effect sizes of at least f=0.25 (medium effect size).

### Experimental procedure

For the Pilot study eligibility was assessed using a structured telephone interview. For the Case-control study, participants attended a screening session, completing the Mini International Neuropsychiatric Interview (Sheehan et al., 1997), the Family Interview for Genetic Studies (Maxwell, 1992) and symptom questionnaires before returning for cognitive testing in a separate session.

In both studies, participants completed the digit span forwards and backwards (Wechsler, 1997), the Wechsler Test of Adult Reading (WTAR(Holdnack, 2001), the Apathy Evaluation Scale (AES(Marin, Biedrzycki, & Firinciogullari, 1991), the Beck Depression Inventory (BDI-II(Beck, Steer, & Brown, 1996), the Dysfunctional Attitudes Scale (DAS-SF1-2(Beevers et al., 2007), the Life Orientation Test-Revisited (LOTR(Scheier, Carver, & Bridges, 1994), the Snaith-Hamilton Pleasure Scale (SHAPS(Snaith et al., 1995), the State-Trait Anxiety Inventory for Adults (STAI(Spielberger, 1970), and the Temporal Experience of Pleasure Scale (TEPS(Gard, Gard, Kring, & John, 2006). In the Pilot study, participants completed the Chapman Physical Anhedonia Scale (CPAS(Chapman, Chapman, & Raulin, 1976). In the Case-control study, participants completed the Hamilton Depression Rating Scale (HAM-D(Hamilton, 1967).

### Apple Gathering Task (AGT)

The AGT measures willingness to exert physical effort for reward (Chong et al., 2015). Before the main task, participants performed a six-trial calibration phase, squeezing a hand-dynamometer as hard as possible with their non-dominant hand to fill an on-screen gauge. The peak force from the final three trials determined their maximum voluntary contraction (MVC), setting comparable effort levels across subjects. Participants then attempted four effort levels (20%, 40%, 60% and 80% of MVC).

On each trial in the main task, participants saw a tree containing a number of apples representing the available reward, and a bar on the tree trunk representing the force required to obtain them (Figure 1A; see SI). There were four levels of reward (3, 6, 9, or 12 apples) and effort (20%, 40%, 60% and 80%). Each reward/effort combination was repeated five times, resulting in a total of 80 trials. Participants could accept or refuse offers. For refused offers, ‘no response required’ was displayed, followed by the next decision. An accepted offer required squeezing the hand-dynamometer at or above the effort level to win points, followed by feedback. The next trial started immediately after the feedback. To mitigate fatigue, the exertion phase was omitted and “No response required” appeared on 25% of accepted trials.

**Figure 1:**
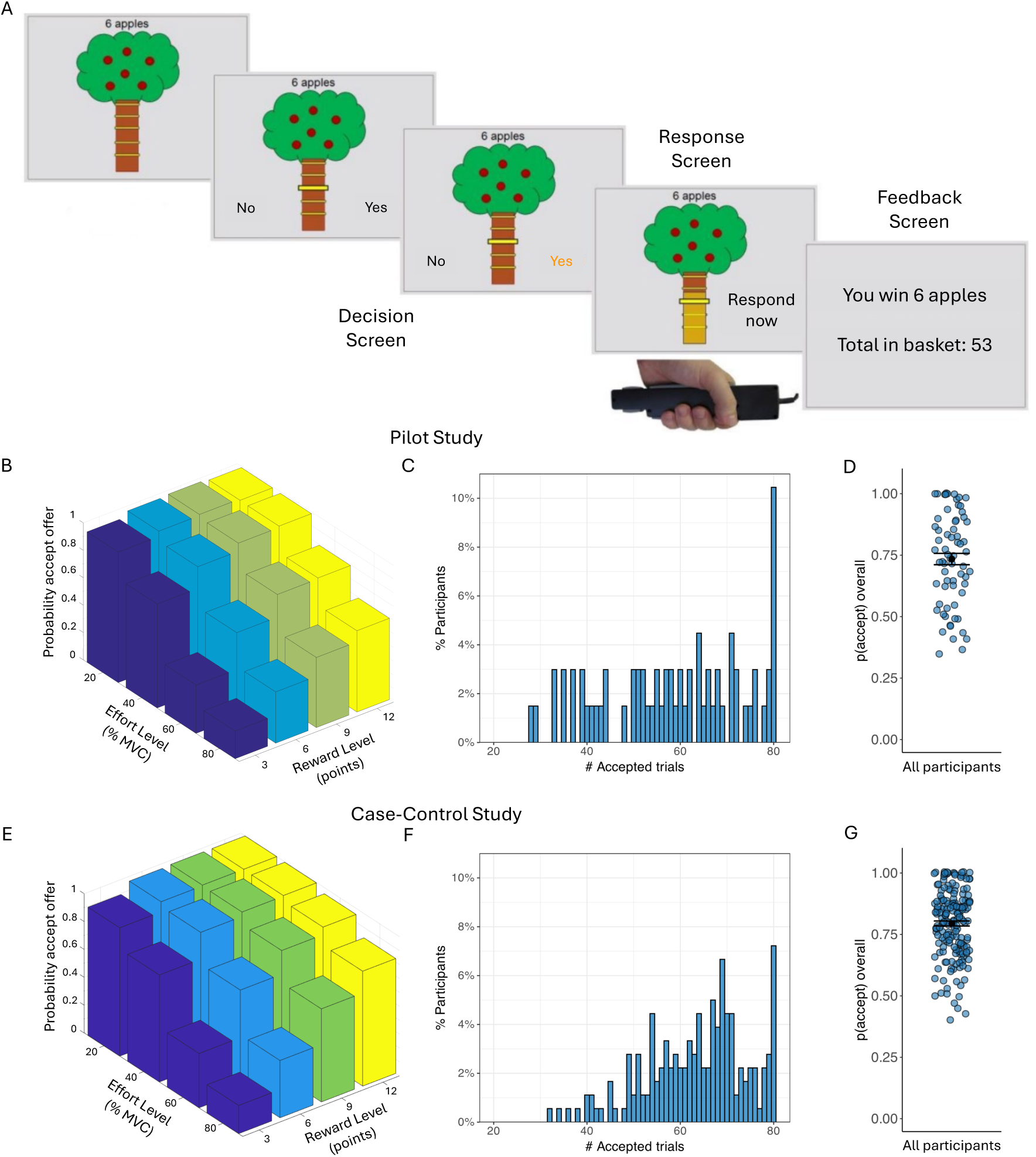
Apple Gathering Task (AGT) and acceptance rates. (A) On each trial, participants are given a different offer comprising a number of apples (3, 6, 9, or 12 apples) for a given effort cost (20%, 40%, 60% or 80% of their maximum grip strength). Participants can either accept the offer or refuse the offer. If the offer is accepted, participants need to squeeze the gripper to the required effort level (or above) for 3 seconds in order to win the apples on this trial. For refused offers, ‘no response required’ was displayed, followed by the next decision. (B) Average acceptance rates as a function of reward level (number of points) and effort level (% MVC) for the Pilot study. (C) Distribution of the number of accepted offers (out of 80) in the Pilot. (D) Overall probability to accept offers for all Pilot participants. Black dots represent the mean and error bars represent the standard error of the mean. (E) Average acceptance rates as a function of reward level and effort level across all groups in the Case-control study. (F) Distribution of the number of accepted offers (out of 80) across all groups in the Case-control study. (G) Overall probability to accept offers for all Case-control participants. Black dots represent the mean and error bars represent the standard error of the mean. Note that raw data is presented but analyses were conducted on arcsine transformed data.

### Statistical Analysis

Repeated-measures analysis of variance (ANOVA) was used to examine how acceptance rates varied as a function of reward and effort. For the Case-control study, group was added as a between-subjects measure. Greenhouse-Geisser correction was applied when sphericity violations occurred. Acceptance rates were arcsine transformed prior to analysis to satisfy Gaussian assumptions. Decision reaction times, success rates and fatigue effects were also analysed (see SI). Covariates in analyses (age, see below) were mean corrected. To examine whether altered effort-based decision making is a trait marker of depression, as opposed to a consequence of experiencing current symptoms, we conducted two planned comparisons following the initial ANOVA: 1) MDD+REM vs REL+CTR; 2) MDD+REM+REL vs CTR.

Questionnaire data was synthesized using exploratory factor analysis on the total score of each questionnaire using Promax rotation (see SI).

### Computational Analysis

Computational analysis was performed to compare competing hypotheses of reward and effort contributions to decision-making, and to estimate participant-level parameters. All models were implemented using hierarchical Bayesian estimation in Stan (Carpenter et al., 2017). Participants in the Pilot study were fit under the same group-level priors. For the Case-control study, all participants were fit under the same prior for parameters of no interest and separate group-level priors were used for the parameter of interest (one parameter at a time; see SI; Figure S1). Sensitivity analyses with separate group priors did not materially impact the estimated parameters or group comparison results (Figure S2).

### Winning model

Seventy models of varying complexity were compared to select the winning model (see SI). The winning model in both studies was a four-parameter model including two effort sensitivity terms (linear: LinE - more negative=greater subjective effort, and quadratic: E^2^ – effort cost increases disproportionately with increasing effort when E^2^<0), a linear reward sensitivity term (LinR; more positive=greater subjective reward), and an acceptance bias term (K) representing the overall tendency to accept offers independent of reward or effort (higher=more likely to accept; Figure S1). Unlike the reward and effort sensitivity parameters, which capture how participants evaluate the components of a given offer (e.g., how much value they assign to a reward, or how aversive they evaluate increasing effort to be), the K parameter captures an overall motivation to exert effort, irrespective of the potential reward or exertion required (Figure S1D). In the Case-control study, owing to high trade-off between the LinE and E^2^ parameters and limited LinE range, the LinE term was constrained to that of the average value estimated from the Pilot study (ConstE=-15; see SI). Together these parameters could accurately recapitulate both the group-level pattern of choices and individual differences in acceptance rates (Figure S3; Pilot: r=0.997; Case-control: r=0.999). All parameters showed high recoverability (Figure S4).

## Results

Participant characteristics are presented in Table 1.

**Table 1:**
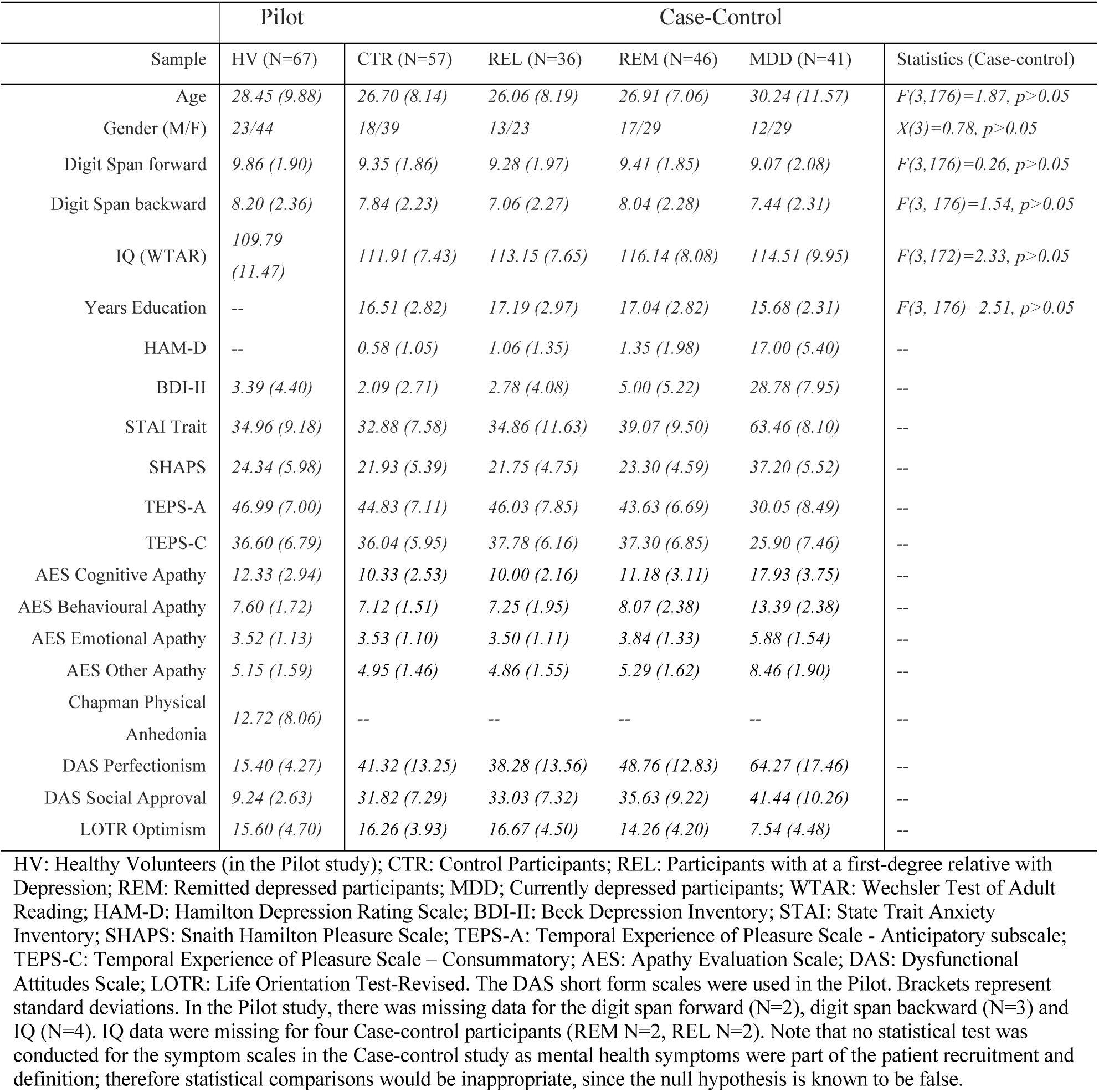
Participant characteristics.

### Model-agnostic AGT analysis

In preliminary analyses we found that older individuals accepted more offers overall (Pearson r=0.20, p=0.014), and therefore age was included as a covariate in all Case-control analyses. We observed wide variability in overall acceptance rates, ranging from 35-100% in the Pilot and 40-100% in the Case-control study (Figure 1C&E). As expected, acceptance rates decreased significantly as effort increased (Pilot: F(1.57,103.77)=120.19, p<0.001; Case-control: F(1.75,306.31)=351.79, p<0.001), and increased significantly as reward increased (Pilot: F(1.63,107.57)=92.02, p<0.001; Case-control: F(1.65,288.06)= 325.45, p<0.001). There was a significant reward-by-effort interaction in both studies, driven by particularly high acceptance rates for high-reward/low-effort trials (Pilot: F(4.25,280.88)=16.42, p<0.001; Case-control: F(4.95,866.79)=90.13, p<0.001; Figure 1B&D).

In the Case-control study we detected a significant main effect of group (F(3,175)=3.26, p=0.023; Figure 2), but all interactions with group were non-significant. Post-hoc tests revealed significantly lower acceptance rates for the REM and MDD groups compared to REL (REM vs REL: Cohen’s d=0.63, p=0.007; MDD vs REL: d=0.41, p=0.015; reported effect sizes exclude covariates). Planned comparisons revealed that the combined MDD+REM group accepted significantly lower acceptance rates than the combined REL+CTR group (d=0.38, p=0.004), but the difference between the combined MDD+REM+REL group and the CTR group was non-significant (d=0.17, p=0.22). Reported effect sizes exclude covariates.

**Figure 2:**
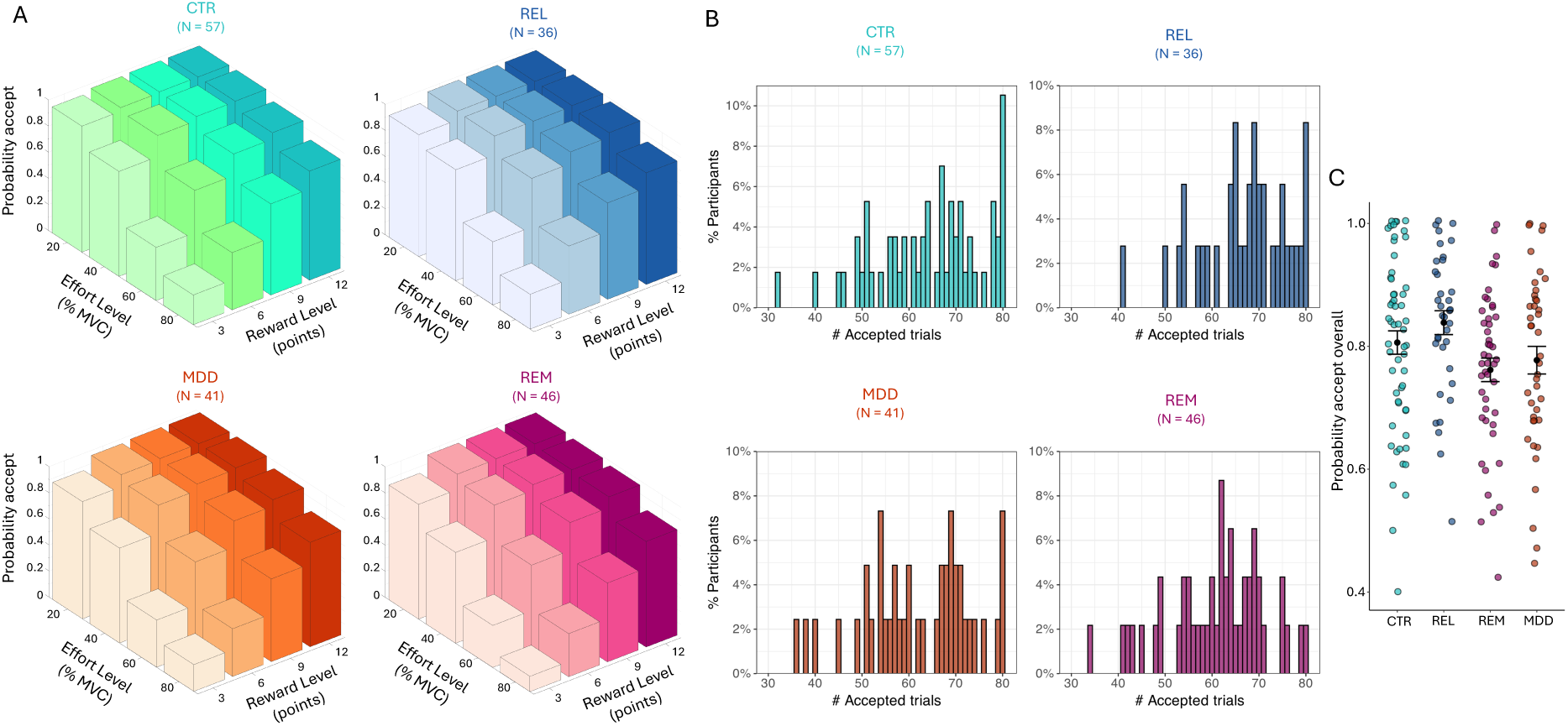
Acceptance rates for the Case-control study. (A) Average acceptance rate as a function of reward level (points) and effort level (% MVC) for the control (CTR), first degree relatives (REL), patients with current depression (MDD), and remitted depression (REM) group. (B) Distribution of the number of accepted offers for each group. (C) Overall probability to accept offers for each group. Black dots represent the mean and error bars represent the standard error of the mean. Note that raw data is presented but analyses were conducted on arcsine transformed data.

There were no significant group differences in success rates, decision RTs, MVCs or either immediate or cumulative fatigue (all p>0.05, see SI; Figure S5&S6). Importantly, success rate at the highest effort level was above 80% across all groups.

### Questionnaire factor analysis

Factor analysis was performed on questionnaire measures for both studies. Despite minor differences in measures (CPAS and the short form of the DAS was only included in the Pilot; HAM-D was only included in the Case-control study), a similar four-factor solution was obtained in both studies (Table S1). The factors were (named according to the scales displaying the highest loadings (>0.3)): “Low-mood”, with loadings for BDI-II, HAM-D, LOTR and STAI; “Apathy”, mostly comprising the AES sub-scales; “Hedonia”, including the SHAPS and TEPS (owing to scoring conventions these load with opposite signs), and (in the Pilot) the CPAS; and “Dysfunctional Attitudes”, comprising the two DAS sub-scales.

### Computational AGT analysis

#### Model comparison

The winning model in both studies was a derivative of a four-parameter model including an acceptance bias parameter (K), linear reward (LinR) and effort (LinE) sensitivity parameters, and a quadratic effort parameter (E^2^; Figure 3; Figures S7&8). In the Case-control study, the LinE term was set at ConstE=-15 (see Materials and Methods and SI).

**Figure 3:**
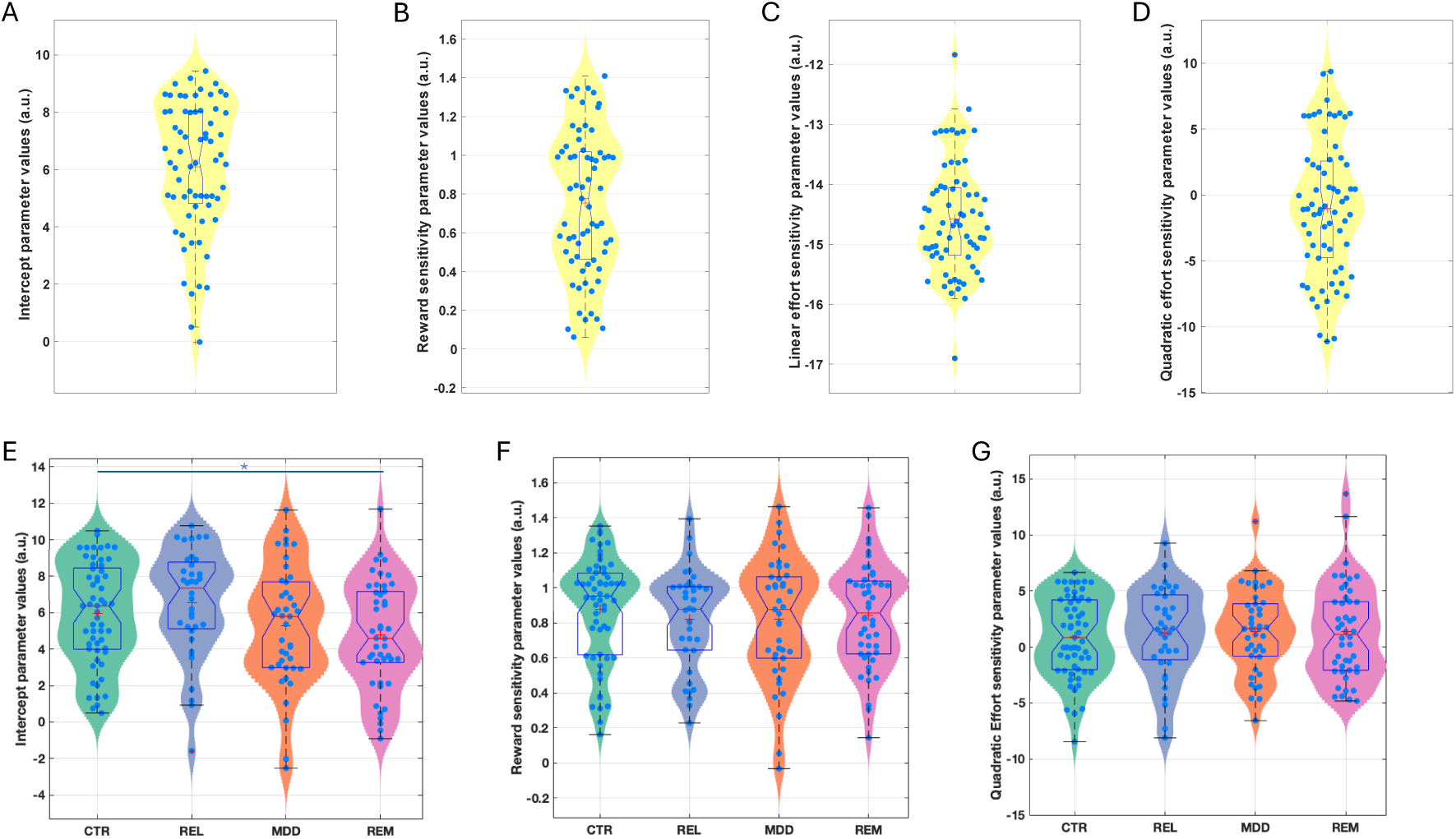
Estimated model parameters for the Pilot (A-D) and Case-control (E-G) study. Figures are showing violin and boxplots as well as the mean (plus sign) and median (notch) for (A) estimated intercept/bias (K), (B) reward sensitivity (LinR), (C) linear effort (LinE), and (D) quadratic effort sensitivity (E^2^) parameter values from the winning model in the pilot study. Data are shown for estimated (E) intercept/bias (K), (F) reward sensitivity (LinR), and (G) quadratic effort (E^2^) sensitivity parameter values from the winning model in the Case-control study. CTR: Control group; REL: First-degree relative group, MDD: Current depression group; REM: Remitted depression group. *Denotes significance at p<0.05.

#### Case-control comparisons

Participant-level parameters were compared between the groups including age as a covariate (Figure 3). The K (acceptance bias) parameter differed significantly between the groups (F(3,175)=3.19, p=0.025). Post-hoc tests revealed that this was driven by the REM (d=0.61, p=0.006) and MDD (d=0.41, p=0.032) groups having a lower acceptance bias than the REL group, and the REM group having a lower acceptance bias than the HC group (d=0.41, p=0.04). Planned comparisons revealed that the combined REM+MDD group had a lower acceptance bias than the combined REL+CTR group (d=0.39, F(1,177)=8.26, p=0.005). However, the combined REL+REM+MDD group did not differ from the CTR group (d=0.16, p=0.276). No other parameter differed significantly between groups (LinR: F(3,175)=0.33, p=0.806; E^2^: F(3,175)=0.13, p=0.942). Reported effect sizes exclude covariates.

The above analysis suggests that lower acceptance rates in the depression groups were driven by a lower overall willingness to exert effort, not alterations in reward or effort sensitivity.

### Associations between parameters and symptoms

#### Pilot study

In the Pilot study, LinR correlated negatively with Low Mood (r=-0.344, p=0.004; Figure 4), suggesting that participants with greater anxiety/depression perceived rewards as less valuable. The E^2^ parameter correlated negatively with Low Mood (r=-0.25, p=0.04) and positively with Hedonia (r=0.248, p=0.043), suggesting that participants with greater anxiety/depression and anhedonia perceived increasing levels of effort as disproportionately costly.

**Figure 4:**
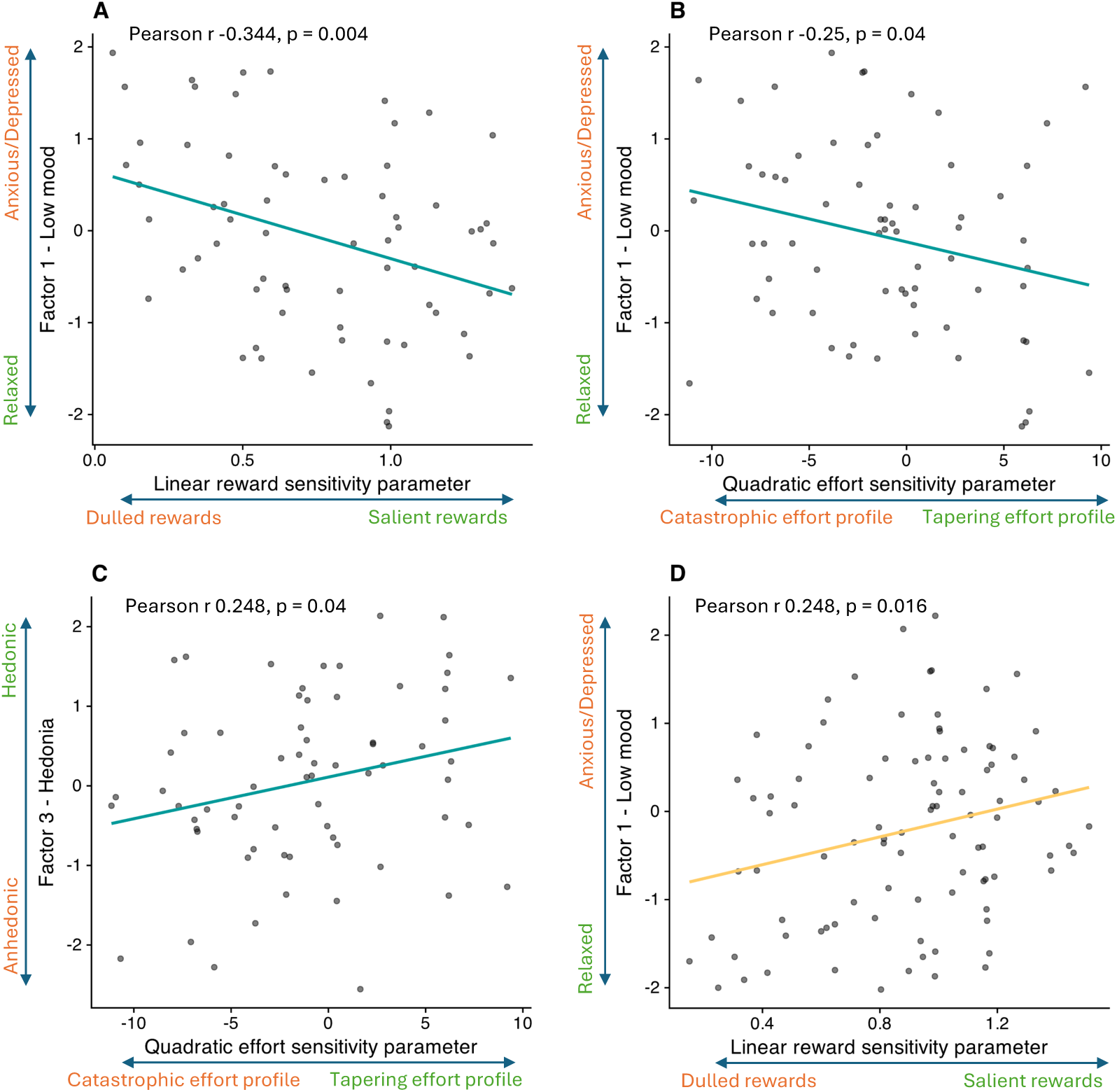
Correlations between computational parameters and symptom factors. (A) Correlation between the Low Mood factor and the linear reward sensitivity (LinR) parameter in the pilot study. (B) Correlation between the Low Mood factor and the quadratic effort sensitivity (E^2^) parameter in the pilot study. (C) Correlation between the Hedonia factor and the quadratic effort sensitivity (E^2^) parameter in the pilot study. (D) Correlation between the Low Mood factor and the linear reward sensitivity (LinR) parameter in the case-control study for the CTR+REL group only.

#### Case-control study

In the Case-control study, no significant associations between symptom factors and computational parameters were observed, either in the combined CTR+REL+REM sample or in the MDD group alone. In the combined CTR+REL group, surprisingly (and in contrast to the corresponding result in the Pilot study) LinR correlated with Low Mood (r=0.248, p=0.016; Figure 4).

## Discussion

Impaired motivation is a hallmark of depression, but the underlying cognitive and computational processes are poorly understood. Consistent with our hypotheses, we found that depression is characterised by a lower willingness to engage in effort. Our computational analyses suggest that this difference was not driven by altered effort or reward sensitivity, but rather by a lower overall tendency to avoid exerting effort, reflected in the acceptance bias parameter. The presence of lower acceptance bias in both the remitted and currently depressed groups suggests that this decision-making process could represent a trait-like feature, as proposed by neurocognitive models of depression (Eshel & Roiser, 2010; Pizzagalli, 2022; Pringle et al., 2013; Roiser et al., 2012), or possibly a “scar” effect of having previously experienced depression (Rohde, Lewinsohn, & Seeley, 1990).

These findings align with previous work demonstrating lower willingness to exert effort for reward in depression (Treadway, Bossaller, Shelton, & Zald, 2012; Treadway et al., 2009; Yang et al., 2016; Yang et al., 2014; Zou et al., 2020) and following recovery (Kuhn et al., 2025). Note that this last study also identified a counter-intuitive greater sensitivity to high rewards in remitted patients which we did not observe here. Interestingly, our findings challenge a key assumption that mis-calibrated reward or effort valuation drives lower willingness to exert effort (Husain & Roiser, 2018; Pessiglione et al., 2018; Treadway & Zald, 2011). By examining a large space of computational models, we instead suggest that a different process is at play: acceptance bias. While surprising and not entirely straightforward to interpret, this result clarifies prior findings from tasks that were unable to disambiguate between these factors. One possibility is that lower acceptance bias could be driven by lower confidence in being able to achieve the required effort, despite individual calibration. This would indicate that metacognitive processes might play a role in motivational impairment. Indeed, previous studies have identified that transdiagnostic symptoms of apathy, poor self-esteem and low mood are associated with low confidence in perceptual decisions (Moses-Payne, Rollwage, Fleming, & Roiser, 2019; Rouault, Seow, Gillan, & Fleming, 2018), independent of accuracy, mirroring our findings. This could plausibly be related to low global expectations of success in depression (Huys, Daw, & Dayan, 2015; Seow, Rouault, Gillan, & Fleming, 2021; Stephan et al., 2016). It is also possible that fatigue may influence acceptance bias, although we did not detect any group differences in sensitivity analyses examining task-related fatigue.

An important goal of neuroscientific research in depression is to determine whether cognitive disruptions are simply a consequence of depressive symptoms, or whether they contribute causally to their development (Halahakoon, Lewis, & Roiser, 2019). We observed a similar pattern of effort-based decisions for reward in both current and remitted depressed groups. This suggests that a general bias against exerting effort might represent a core feature of depression. However, we did not observe a similar pattern in never-depressed first-degree relatives. Therefore, an alternative interpretation might be that lower willingness to exert effort is a consequence rather than a cause of depression, which does not recover after symptoms remit (akin to a ‘scar’ effect). A third possibility is that the first-degree relatives we tested were not at elevated risk for depression, but were instead actually resilient: depression often presents early in life and our REL sample was older (mean age∼26) than the peak onset of depressive symptoms (Lewinsohn, Clarke, Seeley, & Rohde, 1994; Solmi et al., 2022). Thus, many individuals in this group might never go on to develop depression. Future studies should recruit younger samples and test these predictions longitudinally.

If confirmed as a risk factor for depression, lower overall willingness to exert effort may represent a fruitful target for intervention. Such targets are sorely needed for motivational symptoms, which are particularly difficult to treat and constitute an area of unmet clinical need (McMakin et al., 2012; Uher et al., 2012). Treatments that can boost engagement in activities through altering effort acceptance bias could be particularly effective in treating or even preventing depression. Elements of behavioural activation therapy, such as activity scheduling, might play such a role, as these require acting on pre-planned commitments rather than internal states (Martell, Dimidjian, & Herman-Dunn, 2013). Interestingly, this component has recently been shown to alter effort processing in healthy participants, offering proof-of-concept of this idea (Norbury, Hauser, Fleming, Dolan, & Huys, 2024).

### Limitations

While our model of effort-based decision-making replicated across studies, relationships with specific symptom factors were inconsistent, potentially owing to the limited sample size of the Case-control study. Care should also be taken in interpreting these associations as they were exploratory and would not survive corrections for multiple comparisons. Larger studies are needed to confirm these associations. An important strength of the effort task we employed is its individually tailored effort levels, which resulted in >80% average success rates at all effort levels. Nonetheless, it remains possible that occasional failures to obtain reward following exertion could have influenced decisions in some participants. Importantly, however, there was no group difference in success rates—or in effort sensitivity—making it unlikely that the lower effort acceptance bias we observed in depression was influenced by success rates. Additionally, fourteen participants had to be excluded as they were unable to achieve the highest effort level, likely due to issues with calibrating the grip squeeze device. Finally, we interpreted differences in the acceptance bias parameter as reflecting an overall lower willingness to exert effort. However, it is possible that this difference is not specific to effort costs but instead reflects a broader disruption in cost-benefit decision-making – we cannot test this possibility directly here, as our study design only included decisions relating to effort. That said, previous research has not identified differences in other types of reward discounting in depression, for example willingness to take risks (probability discounting; (Charpentier, Aylward, Roiser, & Robinson, 2017), although there is some evidence of elevated delay aversion (temporal discounting), at least in currently symptomatic individuals (Pulcu et al., 2014).

### Conclusion

This study advances our understanding of motivational impairment in depression by identifying low effort acceptance bias, independent of reward or effort sensitivity, as a computational feature of both current and remitted depression. These results raise the possibility that interventions that boost this bias could improve treatment outcomes.

## Ethical standards

The authors assert that all procedures contributing to this work comply with the ethical standards of the relevant national and institutional committees on human experimentation and with the Helsinki Declaration of 1975, as revised in 2008.

## References

Adams, R. A., Huys, Q. J., & Roiser, J. P. (2016). Computational Psychiatry: towards a mathematically informed understanding of mental illness. J Neurol Neurosurg Psychiatry, 87(1), 53–63. doi:10.1136/jnnp-2015-310737

Amsterdam, J. D., Settle, R. G., Doty, R. L., Abelman, E., & Winokur, A. (1987). Taste and smell perception in depression. Biol Psychiatry, 22(12), 1481–1485. doi:10.1016/0006-3223(87)90108-9

Beck, Aaron T, Steer, Robert A, & Brown, Gregory K. (1996). Beck depression inventory.

Beevers, C. G., Strong, D. R., Meyer, B., Pilkonis, P. A., & Miller, I. R. (2007). Efficiently assessing negative cognition in depression: an item response theory analysis of the Dysfunctional Attitude Scale. Psychol Assess, 19(2), 199–209. doi:10.1037/1040-3590.19.2.199

Berwian, I. M., Wenzel, J. G., Collins, A. G. E., Seifritz, E., Stephan, K. E., Walter, H., & Huys, Q. J. M. (2020). Computational Mechanisms of Effort and Reward Decisions in Patients With Depression and Their Association With Relapse After Antidepressant Discontinuation. JAMA Psychiatry, 77(5), 513–522. doi:10.1001/jamapsychiatry.2019.4971

Bonnelle, V., Manohar, S., Behrens, T., & Husain, M. (2016). Individual Differences in Premotor Brain Systems Underlie Behavioral Apathy. Cereb Cortex, 26(2), 807–819. doi:10.1093/cercor/bhv247

Bonnelle, V., Veromann, K. R., Burnett Heyes, S., Lo Sterzo, E., Manohar, S., & Husain, M. (2015). Characterization of reward and effort mechanisms in apathy. J Physiol Paris, 109(1-3), 16–26. doi:10.1016/j.jphysparis.2014.04.002

Bylsma, L. M., Taylor-Clift, A., & Rottenberg, J. (2011). Emotional reactivity to daily events in major and minor depression. J Abnorm Psychol, 120(1), 155–167. doi:10.1037/a0021662

Carpenter, B., Gelman, A., Hoffman, M. D., Lee, D., Goodrich, B., Betancourt, M., … Riddell, A. (2017). Stan: A Probabilistic Programming Language. J Stat Softw, 76. doi:10.18637/jss.v076.i01

Chapman, L. J., Chapman, J. P., & Raulin, M. L. (1976). Scales for physical and social anhedonia. J Abnorm Psychol, 85(4), 374–382. doi:10.1037//0021-843x.85.4.374

Charpentier, C. J., Aylward, J., Roiser, J. P., & Robinson, O. J. (2017). Enhanced Risk Aversion, But Not Loss Aversion, in Unmedicated Pathological Anxiety. Biological Psychiatry, 81(12), 1014–1022. 10.1016/j.biopsych.2016.12.010

Chentsova-Dutton, Y., & Hanley, K. (2010). The effects of anhedonia and depression on hedonic responses. Psychiatry Res, 179(2), 176–180. doi:10.1016/j.psychres.2009.06.013

Chong, T. T., Apps, M., Giehl, K., Sillence, A., Grima, L. L., & Husain, M. (2017). Neurocomputational mechanisms underlying subjective valuation of effort costs. PLoS Biol, 15(2), e1002598. doi:10.1371/journal.pbio.1002598

Chong, T. T., Bonnelle, V., Manohar, S., Veromann, K. R., Muhammed, K., Tofaris, G. K., … Husain, M. (2015). Dopamine enhances willingness to exert effort for reward in Parkinson’s disease. Cortex, 69, 40–46. doi:10.1016/j.cortex.2015.04.003

Cléry-Melin, M. L., Schmidt, L., Lafargue, G., Baup, N., Fossati, P., & Pessiglione, M. (2011). Why don’t you try harder? An investigation of effort production in major depression. PLoS One, 6(8), e23178. doi:10.1371/journal.pone.0023178

Costello, H., Husain, M., & Roiser, J. P. (2024). Apathy and Motivation: Biological Basis and Drug Treatment. Annu Rev Pharmacol Toxicol, 64, 313–338. doi:10.1146/annurev-pharmtox-022423-014645

Croxson, P. L., Walton, M. E., O’Reilly, J. X., Behrens, T. E., & Rushworth, M. F. (2009). Effort-based cost-benefit valuation and the human brain. J Neurosci, 29(14), 4531–4541. doi:10.1523/jneurosci.4515-08.2009

Dichter, G. S., Smoski, M. J., Kampov-Polevoy, A. B., Gallop, R., & Garbutt, J. C. (2010). Unipolar depression does not moderate responses to the Sweet Taste Test. Depress Anxiety, 27(9), 859–863. doi:10.1002/da.20690

Draper, A., Koch, R. M., van der Meer, J. W., Aj Apps, M., Pickkers, P., Husain, M., & van der Schaaf, M. E. (2018). Effort but not Reward Sensitivity is Altered by Acute Sickness Induced by Experimental Endotoxemia in Humans. Neuropsychopharmacology, 43(5), 1107–1118. doi:10.1038/npp.2017.231

Eshel, N., & Roiser, J. P. (2010). Reward and punishment processing in depression. Biol Psychiatry, 68(2), 118–124. doi:10.1016/j.biopsych.2010.01.027

Fervaha, Gagan, Foussias, George, Takeuchi, Hiroyoshi, Agid, Ofer, & Remington, Gary. (2016). Motivational deficits in major depressive disorder: Cross-sectional and longitudinal relationships with functional impairment and subjective well-being. Comprehensive Psychiatry, 66, 31–38. 10.1016/j.comppsych.2015.12.004

Gard, David E, Gard, Marja Germans, Kring, Ann M, & John, Oliver P. (2006). Anticipatory and consummatory components of the experience of pleasure: a scale development study. Journal of research in personality, 40(6), 1086–1102.

Halahakoon, D. C., Kieslich, K., O’Driscoll, C., Nair, A., Lewis, G., & Roiser, J. P. (2020). Reward-Processing Behavior in Depressed Participants Relative to Healthy Volunteers: A Systematic Review and Meta-analysis. JAMA Psychiatry, 77(12), 1286–1295. doi:10.1001/jamapsychiatry.2020.2139

Halahakoon, D. C., Lewis, G., & Roiser, J. P. (2019). Cognitive Impairment and Depression-Cause, Consequence, or Coincidence? JAMA Psychiatry, 76(3), 239–240. doi:10.1001/jamapsychiatry.2018.3631

Hamilton, M. (1967). Development of a rating scale for primary depressive illness. Br J Soc Clin Psychol, 6(4), 278–296. doi:10.1111/j.2044-8260.1967.tb00530.x

Holdnack, HA. (2001). Wechsler test of adult reading: WTAR. San Antonio, TX: The Psychological Corporation.

Husain, M., & Roiser, J. P. (2018). Neuroscience of apathy and anhedonia: a transdiagnostic approach. Nat Rev Neurosci, 19(8), 470–484. doi:10.1038/s41583-018-0029-9

Huys, Quentin J.M., Daw, Nathaniel D., & Dayan, Peter. (2015). Depression: A Decision-Theoretic Analysis. Annual Review of Neuroscience, 38(Volume 38, 2015), 1–23. 10.1146/annurev-neuro-071714-033928

Karvelis, Povilas, Paulus, Martin P., & Diaconescu, Andreea O. (2023). Individual differences in computational psychiatry: A review of current challenges. Neuroscience & Biobehavioral Reviews, 148, 105137. 10.1016/j.neubiorev.2023.105137

Klein-Flügge, Miriam C., Kennerley, Steven W., Friston, Karl, & Bestmann, Sven. (2016). Neural Signatures of Value Comparison in Human Cingulate Cortex during Decisions Requiring an Effort-Reward Trade-off. The Journal of Neuroscience, 36(39), 10002–10015. doi:10.1523/jneurosci.0292-16.2016

Kuhn, M., Palermo, E. H., Pagnier, G., Blank, J. M., Steinberger, D. C., Long, Y., … Pizzagalli, D. A. (2025). Computational Phenotyping of Effort-Based Decision Making in Unmedicated Adults With Remitted Depression. Biological psychiatry Cognitive neuroscience and neuroimaging, 10(6), 607–615. 10.1016/j.bpsc.2025.02.006

Kurniawan, I. T., Guitart-Masip, M., Dayan, P., & Dolan, R. J. (2013). Effort and valuation in the brain: the effects of anticipation and execution. J Neurosci, 33(14), 6160–6169. doi:10.1523/jneurosci.4777-12.2013

Le Heron, Campbell, Plant, Olivia, Manohar, Sanjay, Ang, Yuen-Siang, Jackson, Matthew, Lennox, Graham, … Husain, Masud. (2018). Distinct effects of apathy and dopamine on effort-based decision-making in Parkinson’s disease. Brain, 141(5), 1455–1469. doi:10.1093/brain/awy110

Lewinsohn, Peter M., Clarke, Gregory N., Seeley, John R., & Rohde, Paul. (1994). Major Depression in Community Adolescents: Age at Onset, Episode Duration, and Time to Recurrence. Journal of the American Academy of Child & Adolescent Psychiatry, 33(6), 809–818. 10.1097/00004583-199407000-00006

Marin, R. S., Biedrzycki, R. C., & Firinciogullari, S. (1991). Reliability and validity of the Apathy Evaluation Scale. Psychiatry Res, 38(2), 143–162. doi:10.1016/0165-1781(91)90040-v

Martell, C. R., Dimidjian, S., & Herman-Dunn, R. (2013). Behavioral Activation for Depression: A Clinician’s Guide: Guilford Press.

Maxwell, M Elizabeth. (1992). Family Interview for Genetic Studies (FIGS): a manual for FIGS. Bethesda, MD: Clinical Neurogenetics Branch, Intramural Research Program, National Institute of Mental Health.

McCabe, C., Mishor, Z., Cowen, P. J., & Harmer, C. J. (2010). Diminished neural processing of aversive and rewarding stimuli during selective serotonin reuptake inhibitor treatment. Biol Psychiatry, 67(5), 439–445. doi:10.1016/j.biopsych.2009.11.001

McMakin, Dana L, Olino, Thomas M, Porta, Giovanna, Dietz, Laura J, Emslie, Graham, Clarke, Gregory, … Birmaher, Boris. (2012). Anhedonia predicts poorer recovery among youth with selective serotonin reuptake inhibitor treatment–resistant depression. Journal of the American Academy of Child & Adolescent Psychiatry, 51(4), 404–411.

Mkrtchian, Anahit, Valton, Vincent, & Roiser, Jonathan P. (2023). Reliability of Decision-Making and Reinforcement Learning Computational Parameters. Computational Psychiatry. doi:10.5334/cpsy.86

Moses-Payne, M. E., Rollwage, M., Fleming, S. M., & Roiser, J. P. (2019). Postdecision Evidence Integration and Depressive Symptoms. Front Psychiatry, 10, 639. doi:10.3389/fpsyt.2019.00639

Norbury, Agnes, Hauser, Tobias U., Fleming, Stephen M., Dolan, Raymond J., & Huys, Quentin J. M. (2024). Different components of cognitive-behavioral therapy affect specific cognitive mechanisms. Science Advances, 10(13), eadk3222. doi:10.1126/sciadv.adk3222

Peeters, F., Nicolson, N. A., Berkhof, J., Delespaul, P., & deVries, M. (2003). Effects of daily events on mood states in major depressive disorder. J Abnorm Psychol, 112, 203–211.

Pelizza, L., & Ferrari, A. (2009). Anhedonia in schizophrenia and major depression: state or trait? Ann Gen Psychiatry, 8, 22. doi:10.1186/1744-859x-8-22

Pessiglione, M., Vinckier, F., Bouret, S., Daunizeau, J., & Le Bouc, R. (2018). Why not try harder? Computational approach to motivation deficits in neuro-psychiatric diseases. Brain, 141(3), 629–650. doi:10.1093/brain/awx278

Pizzagalli, D. A. (2022). Toward a Better Understanding of the Mechanisms and Pathophysiology of Anhedonia: Are We Ready for Translation? Am J Psychiatry, 179(7), 458–469. doi:10.1176/appi.ajp.20220423

Pringle, A., McCabe, C., Cowen, P. J., & Harmer, C. J. (2013). Antidepressant treatment and emotional processing: can we dissociate the roles of serotonin and noradrenaline? J Psychopharmacol, 27(8), 719–731. doi:10.1177/0269881112474523

Pulcu, E., Trotter, P. D., Thomas, E. J., McFarquhar, M., Juhasz, G., Sahakian, B. J., … Elliott, R. (2014). Temporal discounting in major depressive disorder. Psychol Med, 44(9), 1825–1834. doi:10.1017/s0033291713002584

Rohde, P., Lewinsohn, P. M., & Seeley, J. R. (1990). Are people changed by the experience of having an episode of depression? A further test of the scar hypothesis. J Abnorm Psychol, 99(3), 264–271. doi:10.1037//0021-843x.99.3.264

Roiser, J. P., Elliott, R., & Sahakian, B. J. (2012). Cognitive mechanisms of treatment in depression. Neuropsychopharmacology, 37(1), 117–136. doi:10.1038/npp.2011.183

Rouault, M., Seow, T., Gillan, C. M., & Fleming, S. M. (2018). Psychiatric Symptom Dimensions Are Associated With Dissociable Shifts in Metacognition but Not Task Performance. Biol Psychiatry, 84(6), 443–451. doi:10.1016/j.biopsych.2017.12.017

Salamone, J. D., Yohn, S. E., López-Cruz, L., San Miguel, N., & Correa, M. (2016). Activational and effort-related aspects of motivation: neural mechanisms and implications for psychopathology. Brain, 139(Pt 5), 1325–1347. doi:10.1093/brain/aww050

Saleh, Y., Jarratt-Barnham, I., Petitet, P., Fernandez-Egea, E., Manohar, S. G., & Husain, M. (2023). Negative symptoms and cognitive impairment are associated with distinct motivational deficits in treatment resistant schizophrenia. Mol Psychiatry. doi:10.1038/s41380-023-02232-7

Scheier, M. F., Carver, C. S., & Bridges, M. W. (1994). Distinguishing optimism from neuroticism (and trait anxiety, self-mastery, and self-esteem): a reevaluation of the Life Orientation Test. J Pers Soc Psychol, 67(6), 1063–1078. doi:10.1037//0022-3514.67.6.1063

Schmidt, L., d’Arc, B. F., Lafargue, G., Galanaud, D., Czernecki, V., Grabli, D., … Pessiglione, M. (2008). Disconnecting force from money: effects of basal ganglia damage on incentive motivation. Brain, 131(Pt 5), 1303–1310. doi:10.1093/brain/awn045

Seow, Tricia X. F., Rouault, Marion, Gillan, Claire M., & Fleming, Stephen M. (2021). How Local and Global Metacognition Shape Mental Health. Biological Psychiatry, 90(7), 436–446. 10.1016/j.biopsych.2021.05.013

Sheehan, D. V., Lecrubier, Y., Harnett Sheehan, K., Janavs, J., Weiller, E., Keskiner, A., … Dunbar, G. C. (1997). The validity of the Mini International Neuropsychiatric Interview (MINI) according to the SCID-P and its reliability. European Psychiatry, 12(5), 232–241. 10.1016/S0924-9338(97)83297-X

Snaith, R. P., Hamilton, M., Morley, S., Humayan, A., Hargreaves, D., & Trigwell, P. (1995). A scale for the assessment of hedonic tone the Snaith-Hamilton Pleasure Scale. Br J Psychiatry, 167(1), 99–103. doi:10.1192/bjp.167.1.99

Solmi, Marco, Radua, Joaquim, Olivola, Miriam, Croce, Enrico, Soardo, Livia, Salazar de Pablo, Gonzalo, … Fusar-Poli, Paolo. (2022). Age at onset of mental disorders worldwide: large-scale meta-analysis of 192 epidemiological studies. Molecular Psychiatry, 27(1), 281–295. doi:10.1038/s41380-021-01161-7

Spielberger, Charles Donald. (1970). Manual for the State-Trait Anxiety Inventory (self-evaluation questionnaire). (No Title).

Spijker, J., Bijl, R. V., De Graaf, R., & Nolen, W. A. (2001). Determinants of poor 1-year outcome of DSM-III-R major depression in the general population: results of the Netherlands Mental Health Survey and Incidence Study (NEMESIS). Acta Psychiatrica Scandinavica, 2, 122–130. 10.1034/j.1600-0447.2001.103002122.x

Stephan, K. E., Manjaly, Z. M., Mathys, C. D., Weber, L. A., Paliwal, S., Gard, T., … Petzschner, F. H. (2016). Allostatic Self-efficacy: A Metacognitive Theory of Dyshomeostasis-Induced Fatigue and Depression. Front Hum Neurosci, 10, 550. doi:10.3389/fnhum.2016.00550

Treadway, M. T., Bossaller, N. A., Shelton, R. C., & Zald, D. H. (2012). Effort-based decision-making in major depressive disorder: a translational model of motivational anhedonia. J Abnorm Psychol, 121(3), 553–558. doi:10.1037/a0028813

Treadway, M. T., Buckholtz, J. W., Schwartzman, A. N., Lambert, W. E., & Zald, D. H. (2009). Worth the ‘EEfRT’? The effort expenditure for rewards task as an objective measure of motivation and anhedonia. PLoS One, 4(8), e6598. doi:10.1371/journal.pone.0006598

Treadway, M. T., & Zald, D. H. (2011). Reconsidering anhedonia in depression: lessons from translational neuroscience. Neurosci Biobehav Rev, 35(3), 537–555. doi:10.1016/j.neubiorev.2010.06.006

Uher, R., Perlis, R. H., Henigsberg, N., Zobel, A., Rietschel, M., Mors, O., … McGuffin, P. (2012). Depression symptom dimensions as predictors of antidepressant treatment outcome: replicable evidence for interest-activity symptoms. Psychological Medicine, 42(5), 967–980. doi:10.1017/S0033291711001905

Wechsler, D. (1997). WMS-III Administration and Scoring Manual. In. San Antonio, TX: The Psychological Corporation. Harcourt Brace & Co.

Yang, X. H., Huang, J., Lan, Y., Zhu, C. Y., Liu, X. Q., Wang, Y. F., … Chan, R. C. (2016). Diminished caudate and superior temporal gyrus responses to effort-based decision making in patients with first-episode major depressive disorder. Prog Neuropsychopharmacol Biol Psychiatry, 64, 52–59. doi:10.1016/j.pnpbp.2015.07.006

Yang, X. H., Huang, J., Zhu, C. Y., Wang, Y. F., Cheung, E. F., Chan, R. C., & Xie, G. R. (2014). Motivational deficits in effort-based decision making in individuals with subsyndromal depression, first-episode and remitted depression patients. Psychiatry Res, 220(3), 874–882. doi:10.1016/j.psychres.2014.08.056

Yuen, G. S., Bhutani, S., Lucas, B. J., Gunning, F. M., AbdelMalak, B., Seirup, J. K., … Alexopoulos, G. S. (2015). Apathy in late-life depression: common, persistent, and disabling. Am J Geriatr Psychiatry, 23(5), 488–494. doi:10.1016/j.jagp.2014.06.005

Zou, Y. M., Ni, K., Wang, Y. Y., Yu, E. Q., Lui, S. S. Y., Zhou, F. C., … Chan, R. C. K. (2020). Effort-cost computation in a transdiagnostic psychiatric sample: Differences among patients with schizophrenia, bipolar disorder, and major depressive disorder. Psych J, 9(2), 210–222. doi:10.1002/pchj.316

