## Supplementary Materials for "A computational approach to understanding effort-based decision-making in depression"

### Supplemental Information

#### Participants

##### *Pilot study*

Exclusion criteria included any current/past mental health diagnoses (self-report), marijuana use within four weeks, other recreational drug use within one week, or alcohol use within 24hr. In total twelve participants were excluded for the following reasons: incomplete data owing to technical problems; BDI-II score>20, indicating moderate depression; unreliable grip squeeze calibration (indicated by >200% exerted force during the task relative to maximum voluntary contraction (MVC) during calibration). A further 23 participants were excluded from the behavioural analysis due to low success rates at the highest effort levels (p(success)<33% at the 80% effort level), indicating problems with grip-squeeze calibration, leaving a total of N=67.

##### *Case-control study*

Diagnoses were assessed using the Mini International Neuropsychiatric Interview v5 (MINI[1]). Family history of mental illness was assessed using the Family Interview for Genetic Studies (FIGS[2,3]).

MDD participants had to meet criteria for a current major depressive episode (MDE) according to the MINI, with a score≥8 on the Hamilton Depression Rating Scale (HAM-D[4]); REM participants had to meet criteria for a past MDE, with a score≤7 on the HAM-D. REL and CTR participants had no current or past diagnoses, with a score≤7 on the HAM-D, and REL participants had at least one first-degree relative with current/past depression. Additional exclusion criteria for all groups included: any history of neurological disorders; current use of psychotropic medication (including St John's Wort) within the past four weeks (eight weeks for fluoxetine); marijuana use within the past four weeks or other recreational drug use within the past week; alcohol use within the past 24hr; lifetime manic/hypomanic episode; current alcohol/substance use disorder; lifetime severe alcohol/substance use disorder (occurring outside a depressive episode); and lifetime history of any psychotic disorder. Current or past anxiety disorders and severe alcohol/substance use disorder occurring exclusively within a depressive episode were not exclusion criteria for the MDD/REM groups.

Seven participants were excluded for one of the following reasons: incomplete data; inconsistent reporting of symptoms; or unreliable grip squeeze calibration on the effort task (indicated by >200% exerted force during the task relative to MVC during calibration), leaving N=59 CTR, N=37 REL, N=50 REM and N=48 MDD participants. A further 14 participants (one CTR, one REL, four REM, and eight MDD) were excluded from the behavioural analysis due to low success rates at the highest effort levels (p(success)<33% at the 80% effort level), indicating problems with grip-squeeze calibration, leaving a total of N=180: N=57 CTR, N=36 REL, N=46 REM and N=41 MDD participants

In both studies, participants were compensated £30 for completing symptom questionnaires and a task battery including the AGT (data on other tasks will be reported elsewhere). They could earn a bonus up to £20, depending on performance across all tasks. They were recruited through local advertisements, institutional participant databases and local outpatient psychological treatment services, and all provided written informed consent. All participants were aged 18-60 years, unmedicated and native English speakers. The study received ethical approval from the UCL Research Ethics Committee (fMRI/2013/005) and the London Queen Square NHS Research Ethics Committee (for depressed individuals: 10/H0716/2).

#### Additional Task Details

The AGT was implemented in MATLAB R2016b (MathWorks, USA) using Psychtoolbox (v.3.0.14; psychtoolbox.org). A hand-dynamometer was used for effort exertion (SS25LA – BIOPAC Systems, Inc., USA). Participants could either accept or refuse offers by responding 'yes' or 'no' using the keyboard. During the effort response, immediate visual feedback in the form of a red gauge was displayed on the tree trunk so that participants could adapt their exerted force to match the required effort. A feedback screen then informed the participant whether they had succeeded ("You win *X* apples" or "You win *0* apples" for successful and unsuccessful attempts, respectively). At the end of each block, participants were presented with the total number of points gathered on the block. Participants could rest between blocks to avoid fatigue (self-paced). To mitigate fatigue, "No response required" appeared on 25% of accepted trials. Participants were instructed that these skipped effort exertion trials could occur sporadically to prevent fatigue, but that it would not affect their performance bonus.

There was no fixed amount of money to points across participants. The AGT was one of ten tasks in the Pilot study and one of eight tasks in the Case-Control study. At the end of the session, a random number of trials were chosen for each task and a portion of the £20 bonus pot was allocated proportionally to the number of trials selected for that task. The points won on those trials were converted into money. Participants did not know the exact payoff rule but were told that the bonus would be defined from trials selected at random and to earn as much money as possible they needed to perform well at all tasks.

#### Statistical analysis

Raw data was processed using custom scripts in MATLAB (MathWorks, USA). Repeated-measures analysis of variances (ANOVAs) were performed to analyse whether decision reaction times (RTs) and success rates varied as a function of reward magnitude, effort level, or their interaction. Success rates were arcsine transformed prior to analysis, and RTs were log transformed, to satisfy Gaussian assumptions. For the Case-control study, group was added to the ANOVAs as a between-subjects measure.

For the factor analysis, questionnaire scores were transformed to yield Gaussian distributions, then Z-scored prior to factor analysis. The number of factors included in the final factor solution was determined by visual examination of the resulting scree plots and factors yielding eigenvalues>1. Loadings from the emerging factor solution were then applied to each participant's scores to compute factors.

In the Case-control study, some questionnaire scores had bimodal distributions due to high scores in MDD participants. Therefore, factor analysis was performed on data from the other three groups. MDD questionnaire data was transformed and Z-scored using the resulting transformations and normalization constants. This ensured that the factor solution was not biased by the scores in the MDD group. Missing questionnaire data was imputed at the item level using the mean item score in the corresponding group (Pilot 0.33%, Case-control 0.088% imputed items).

#### Computational Analysis

A computational analysis of decisions was performed to parse the contributions of different processes on decision making. A variety of models with varying complexity were built to test and capture the contribution of various cognitive processes on decisions to accept the offer on a given trial. We started by testing simple assumptions, such as whether participants performed randomly (Null model), and then iteratively improved the models until we could capture decision patterns as parsimoniously as possible.

All models were implemented using hierarchical Bayesian estimation in Stan with parameters estimated using Hamiltonian Markov-Chain Monte Carlo sampling. This approach assumes that each group of participants can be described using a population distribution, which improves parameter estimation accuracy, and includes priors over the parameters of this distribution, which acts as soft constraints on likely parameter ranges.

Model comparison was performed to select the winning model (the model that best captured decisions, whilst accounting for increased model complexity). The Widely Applicable Information Criterion (WAIC) scale and K-fold cross validation (K-fold CV) were used to compare model fits[5,6]. For both, a lower value indicates a better model parsimony, and the relative difference between the winning model and other models can be used to establish the relative strength of evidence for each (akin to Bayes factors[7]). We defined differences in model evidence (**Δ**WAIC) as: weak (0-2); positive (2-6); strong (6-10); very strong (>10)[7]. The WAIC is akin to cross-validation, or other approximations such as the Akaike Information Criterion, but is more sensitive, particularly in hierarchical settings[8]. The more computationally demanding K-fold CV was used when diagnostic measures of the WAIC (pareto-k) suggested that the WAIC approximation was sub-optimal.

For each model, two chains were produced with 1000 warm-up iterations and 4000 post-warm-up iterations per chain. Model convergence was ensured through careful analysis of traceplots and monitoring of the Gelman-Rubin statistic (all potential scale reduction factor: R-hat<1.1[9]).

Recommendations for weakly informative priors in hierarchical logistic regression were used[10,11]. For all models, participant-level parameters were normally distributed and drawn from the group-level parameters (e.g., $\theta_{participant} \sim \mathcal{N}(\mu_{group}, \sigma_{group})$; Figure S1), except for the noise term (see Pilot study in Computational results below) which was beta distributed to enforce a proportion of noise level between zero and one: $\theta_{noise} \sim \beta(\mu_{group}, \sigma_{group})$.

The winning model comprised four parameters: two effort sensitivity terms (linear: LinE, and quadratic: $E^2$), a linear reward sensitivity term (LinR), and an acceptance intercept/bias term (K). Together these parameters captured the full range of choice patterns observed across participants. The model operates as follows: for each offer, the effort level is transformed through the effort sensitivity parameters to yield a subjective value of effort (eq.1). The linear effort sensitivity term scales the value of the effort level (more negative=greater subjective effort), and the quadratic term allows the effort profile to either taper off (effort sensitivity$^2$>0), or to increase disproportionately with increasing effort (effort$^2$ sensitivity<0).

$$eq.1 \quad Subjective\ value\ of\ effort = (LinE \times effort) + (E^2 \times effort^2)$$

Reward is transformed through a linear reward sensitivity term to yield a subjective value of reward (eq.2) (more positive=greater subjective reward).

$$eq.2 \quad Subjective\ value\ of\ reward = (LinR \times reward)$$

The subjective values of reward and effort are then combined to form the subjective value of the offer (eq.3).

$$eq.3 \quad Subjective\ value\ of\ offer = Subjective\ value\ of\ reward + Subjective\ value\ of\ effort$$

The subjective value of the offer is passed through an inverted logit link function with a bias parameter (eq.4). This maps the subjective value of the offer to a probability of acceptance, where the bias (or intercept) term shifts the curve by a constant. The bias term therefore represents the overall tendency to accept offers (higher=more likely to accept), independent of reward or effort.

$$eq.4. \quad Accept\ probability = inv\_logit(K + Subjective\ value\ of\ offer)$$

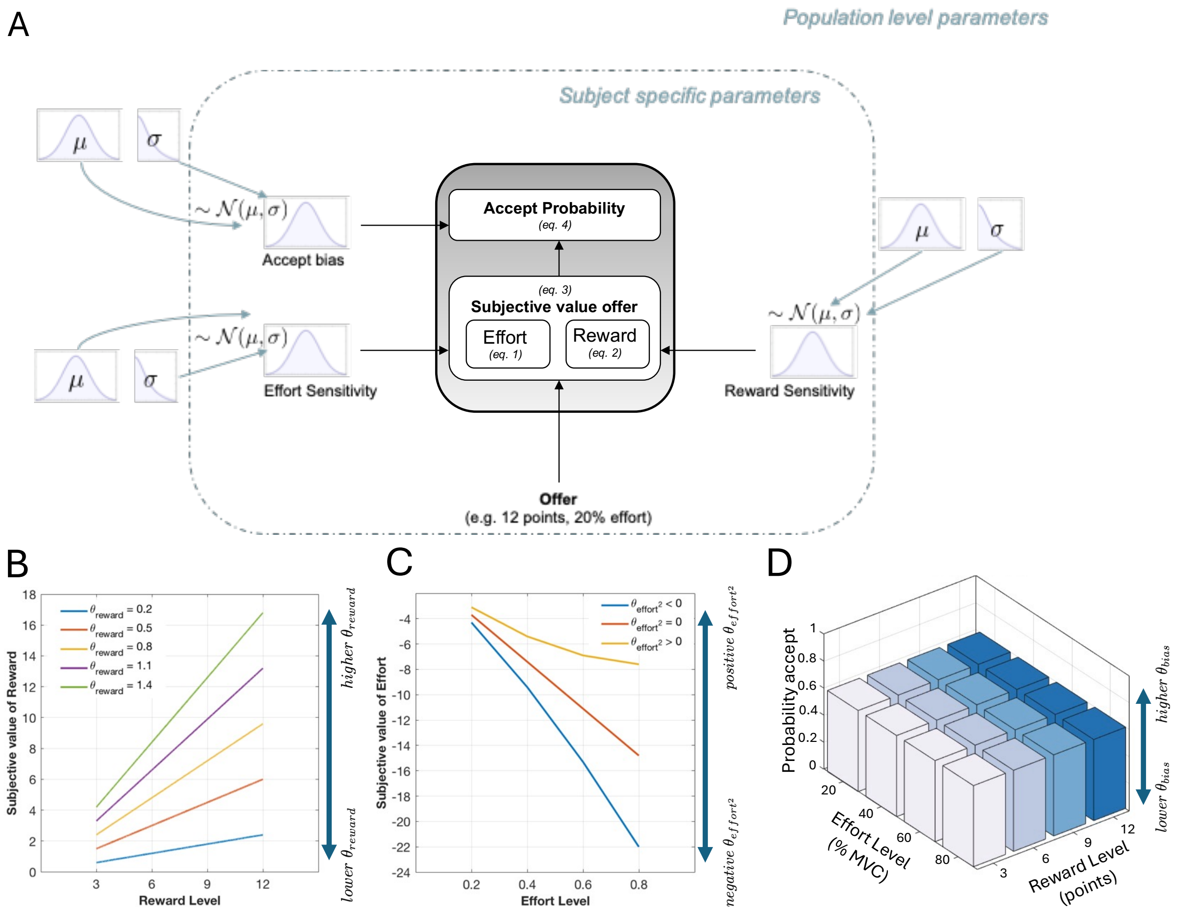


***Figure S1: Graphical depiction of the hierarchical logistic regression model for the Apple Gathering Task.*** *(A) Subject-level parameters are drawn from population level parameters. Subject level parameters transform the true reward and effort levels into subjective values unique to each participant. The subjective value of the offer is passed through an invert logit link function that maps the subjective value into a probability of acceptance. (B) The linear reward sensitivity scales the true rewards such that rewards can be perceived as more or less rewarding than they truly are. (C) Linear and quadratic effort sensitivity terms scale the true effort costs to capture the entire range of possible effort cost profiles. (D) The bias term enables to capture the baseline probability of accepting an offer, regardless of the effort and reward magnitudes.*
*Note that* $\theta_{reward}$ *corresponds to LinR,* $\theta_{effort^2}$ *corresponds to E²* *and* $\theta_{bias}$ *corresponds to K in the equations presented in the main text.*

For the Case-control study, we fitted three separate versions of the winning model, such that all participants were fit under the same prior for parameters of no interest and separate group-level priors were used for the parameter of interest (one for each parameter of interest). In other words, we constructed one version where all participants were fit under the same prior for all parameters but the LinR parameter, which had a separate prior for each group; a second version where participants were fit under the same prior for all but the LinE parameter, which was fit with a separate group-level parameter; and a third version where all participants were fit under the same prior for all but the $E^2$ parameter, which had a group-specific prior. We performed a sensitivity analysis to compare this approach with fitting a model version where all parameters were specified with independent group-level priors. This allowed us to verify that the group-level specifications did not materially impact the estimated parameters or group comparison results (Figure S2).

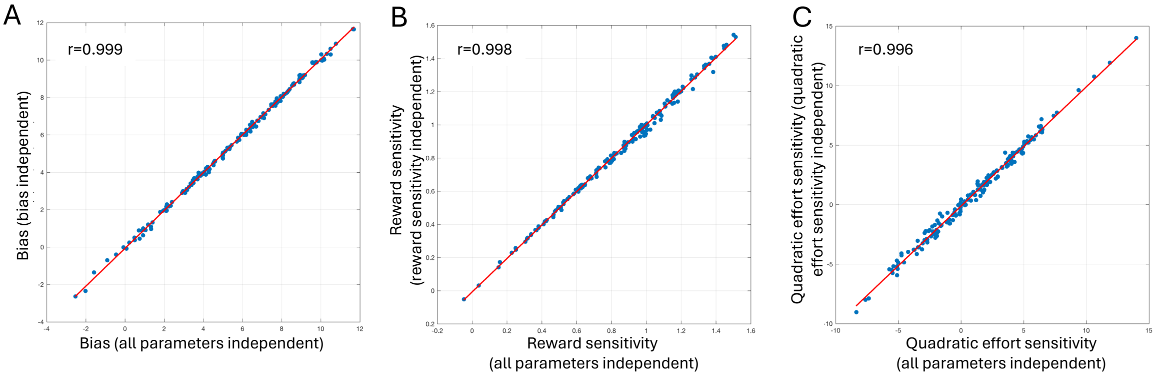


***Figure S2: Sensitivity analysis of inferred parameters with different hierarchical priors.*** *The x-axis is the parameter extracted using independent group-level priors for all parameters, and the y-axis represents the parameter extracted using an independent group-level prior for the (A) bias (K), (B) reward sensitivity (LinR) and (C) quadratic effort sensitivity ($E^2$) parameter with shared group-level priors (pooling) for all other parameters. All parameters were virtually identical with different hierarchical prior setups (r=0.996-0.999).*

#### *Posterior Predictive Checks & Parameter Recovery*

Posterior predictive checks were performed to ascertain that the winning model for each group could adequately recapitulate the observed group-level acceptance rates, as well as recovering inter-individual differences in acceptance rates. Figure S3 shows the predicted pattern of behaviour for each study, as well as the predicted mean overall acceptance rate for each individual against the observed acceptance rate. The winning models in both studies accurately recovered the pattern of behaviour observed in the model-agnostic analysis, as well as almost perfectly recovering individual differences in acceptance rates.

We examined parameter recovery of the winning model by simulating 50 synthetic participants performance on the task. The winning model was then fitted and parameters were extracted for each synthetic participant (all parameters were estimated simultaneously). All parameters were recovered satisfactorily (Pearson's r=0.85-0.99; Absolute Agreement ICC=0.83-0.99; Figure S4).

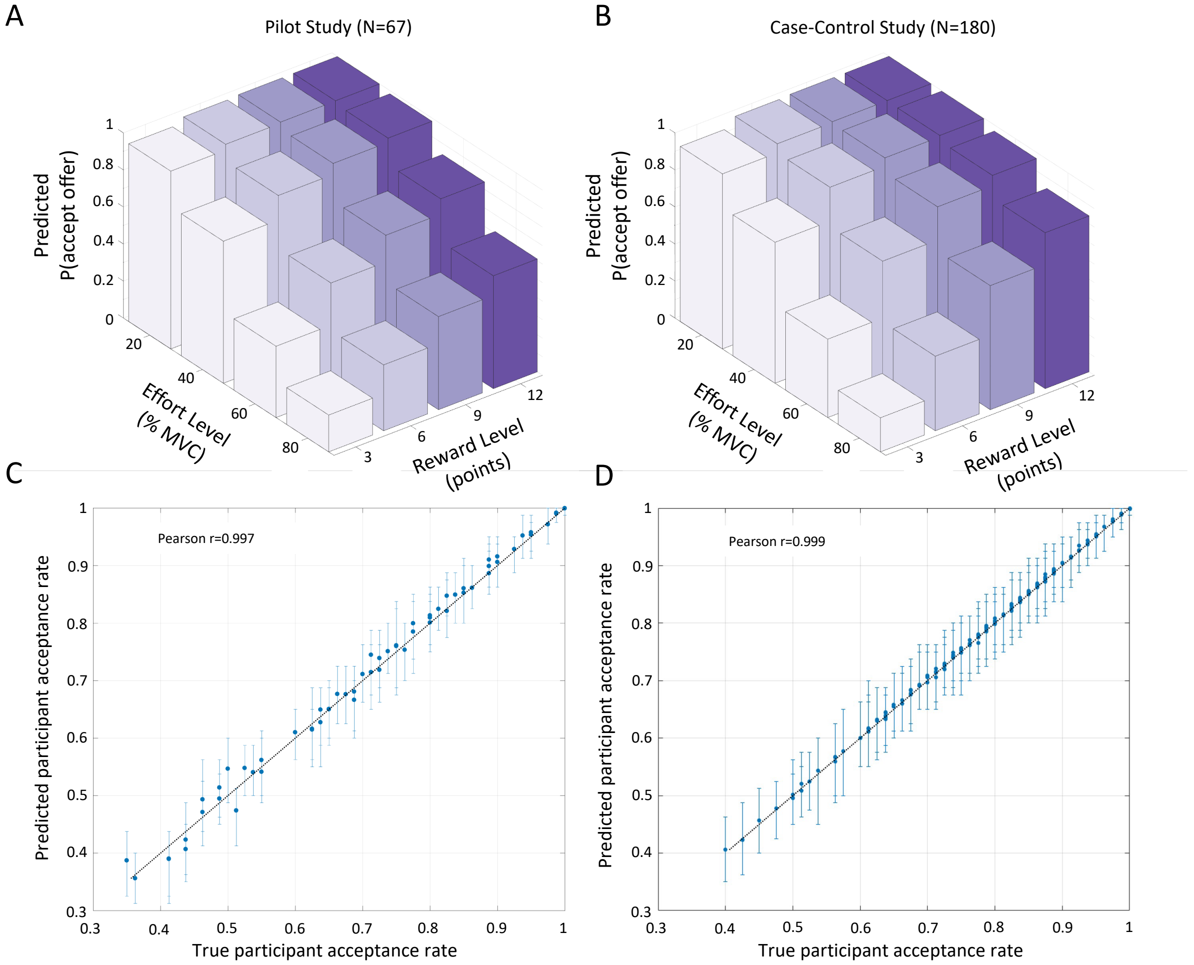


***Figure S3: Posterior predictive checks.*** *(A) Pattern of behaviour predicted from the winning model for the Pilot study and the (B) Case-control study. (C) Inter-individual predicted acceptance rate vs. true participant acceptance rate for the Pilot study and the (D) Case-control study.*

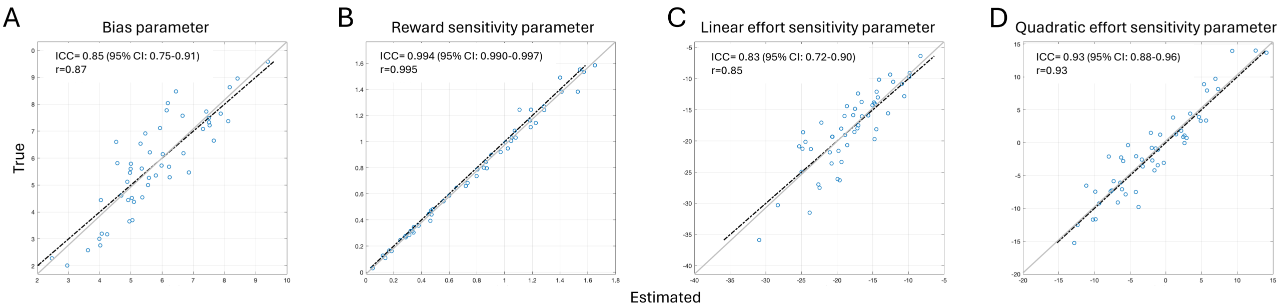


***Figure S4: Parameter recovery of parameters for the winning model****. All parameters (A-D) show high parameter recovery. Parameter recovery is plotted for (A) the bias (K), (B) reward sensitivity (LinR), (C) linear effort sensitivity (LinE) and (D) quadratic effort sensitivity ($E^2$) parameter. ICCs denote absolute agreement, single values. The dotted black line represents line of best fit. The grey line represents a perfect correlation (ICC=1).*

#### Factor analysis

The factor analysis is presented in Table S1.

Table S1: Factor analysis solutions for the Pilot and Case-control (excluding the MDD group) study questionnaire measures.

| | Pilot | | | | Case-Control | | | |
|---|---|---|---|---|---|---|---|---|
| *Estimated Factors* | **LOW MOOD** | **APATHY** | **HEDONIA** | **DYSFUNC. ATTITUDES** | **LOW MOOD** | **APATHY** | **HEDONIA** | **DYSFUNC. ATTITUDES** |
| | **FACTOR 1** | **FACTOR 2** | **FACTOR 3** | **FACTOR 4** | **FACTOR 2** | **FACTOR 1** | **FACTOR 3** | **FACTOR 4** |
| *HAM-D* | -- | -- | -- | -- | **0.60** | -0.01 | 0.00 | -0.20 |
| *BDI-II* | **0.51** | -0.05 | -0.34 | -0.03 | **0.57** | 0.15 | 0.12 | 0.13 |
| *AES Cognitive Apathy* | -0.06 | **0.84** | -0.11 | 0.00 | 0.01 | **0.76** | -0.07 | -0.11 |
| *AES Behavioural Apathy* | 0.25 | **0.66** | 0.15 | -0.09 | 0.15 | **0.67** | 0.04 | -0.02 |
| *AES Emotional Apathy* | -0.24 | **0.59** | -0.21 | -0.02 | -0.19 | **0.66** | -0.11 | 0.04 |
| *AES Other Apathy* | 0.00 | **0.83** | 0.15 | 0.09 | 0.03 | **0.87** | 0.30 | -0.02 |
| *Chapman Physical Anhedonia* | -0.20 | 0.07 | **-0.87** | 0.02 | -- | -- | -- | -- |
| *DAS Perfectionism* | -0.06 | 0.03 | -0.04 | **1.01** | -0.01 | -0.08 | 0.04 | **1.05** |
| *DAS Social Approval* | 0.08 | 0.01 | 0.02 | **0.61** | 0.28 | 0.03 | 0.16 | **0.36** |
| *LOTR Optimism* | **-0.47** | -0.08 | 0.22 | -0.18 | **-0.50** | 0.00 | 0.25 | -0.05 |
| *SHAPS* | 0.25 | 0.01 | **-0.33** | -0.16 | -0.05 | **0.39** | **-0.31** | 0.07 |
| *STAI State* | **0.78** | 0.03 | 0.13 | 0.02 | **0.56** | **0.30** | -0.01 | -0.03 |
| *STAI Trait* | **1.11** | -0.04 | 0.14 | -0.05 | **0.96** | -0.06 | -0.09 | 0.00 |
| *TEPS-A.* | -0.04 | -0.23 | **0.25** | 0.03 | -0.08 | 0.17 | **1.04** | 0.06 |
| *TEPS-C.* | 0.06 | 0.09 | **0.90** | -0.03 | 0.02 | -0.08 | **0.42** | -0.18 |

Questionnaire loadings >0.3 (or highest loading factor) are highlighted in bold. A consistent factor solution was identified between the two studies, other than the order of the first two factors. HAM-D: Hamilton Depression Rating Scale; BDI-II: Beck Depression Inventory; AES: Apathy Evaluation Scale; DAS: Dysfunctional Attitudes Scale; LOTR: Life Orientation Test-Revised; SHAPS: Snaith Hamilton Pleasure Scale; STAI: State Trait Anxiety Inventory; TEPS-A: Temporal Experience of Pleasure Scale - Anticipatory subscale; TEPS-C: Temporal Experience of Pleasure Scale – Consummatory.

#### Success rates and decision response time results

Success rates decreased significantly as effort level increased for both studies (Pilot: $F(2.21,113.15)=24.70$, $p<0.001$; Case-control: $F(3,495)=75.66$, $p<0.001$). Importantly, success rate at the highest effort level was well above 80%, and broadly comparable between the four conditions (Figure S5A-B). In the Case-control study there was no significant effect of group on success rate ($F(3,161)=0.801$, $p>0.05$) or RTs ($F(3,172)=0.464$, $p>0.05$), and all interactions were non-significant.

In the Case-control study, RTs were significantly longer at higher effort levels ($F(2.565,451.49)=87.63$, $p<0.001$), with no main effect of reward ($F(2.875, 506.05)=0.715$, $p>0.05$), and a significant reward-by-effort interaction ($F(8.05, 1418.37)=5.767$, $p<0.001$); this reflects that RTs were particularly long on high-effort/high-reward trials (Figure S5C-D). A similar effect of effort ($F(3, 198)=8.55$, $p<0.001$) on RTs was observed in the Pilot study, with no significant effect of reward ($F(3,198)=0.21$, $p>0.05$), or reward-by-effort interaction ($F(9,594)=1.77$, $p=0.069$).

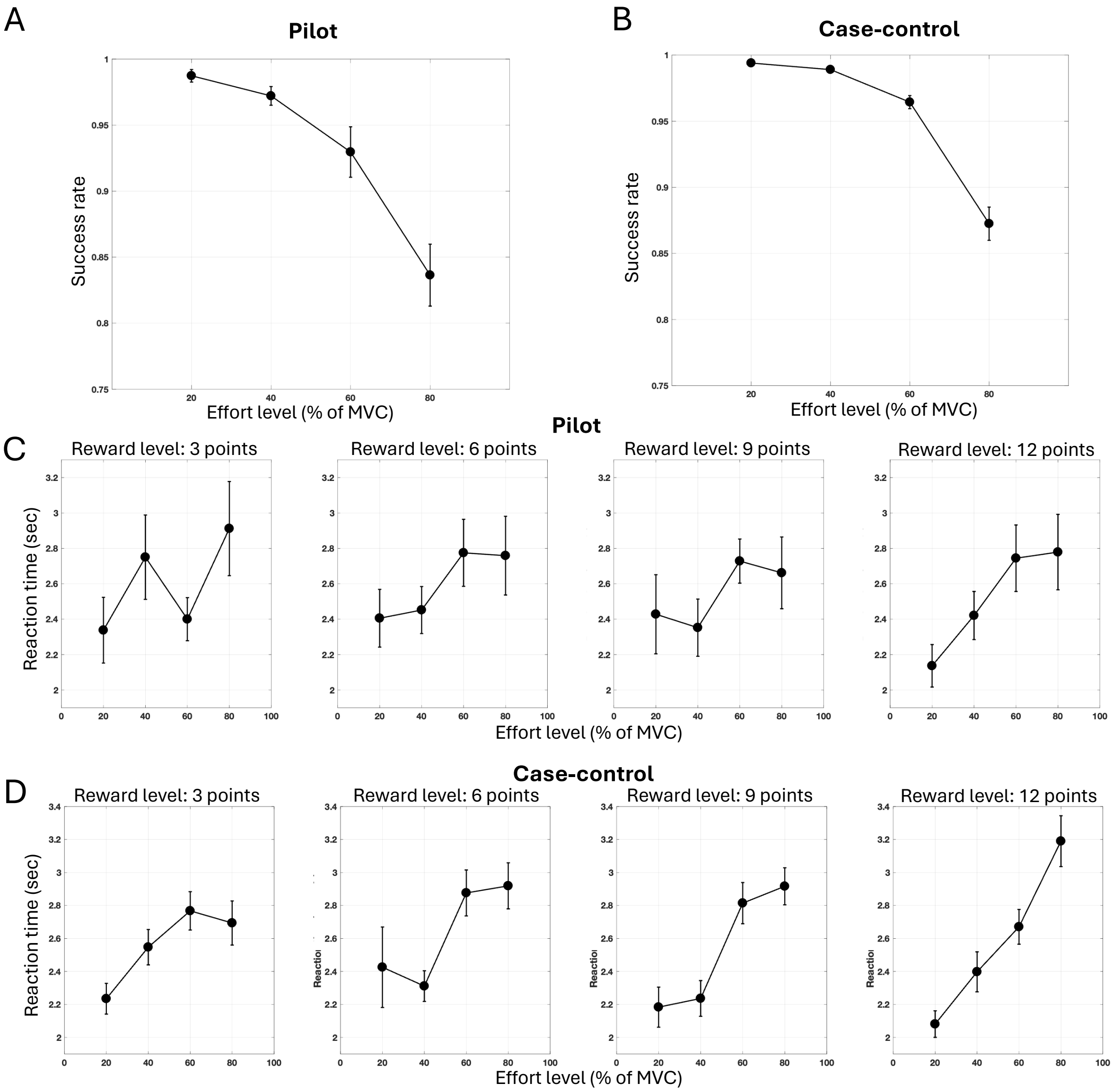


***Figure S5: Success rates and reaction times.*** *(A) Success rates at different effort levels in the Pilot and (B) Case-control study. (C) Decision reaction times at each reward and effort level in the Pilot and (D) Case-control study. Error bars represent standard error of the mean.*

#### Maximum voluntary contraction (MVC) results

There was no significant group effect on MVC in the Case-control study ($F(3,176)=1.83$, $p=0.143$; CTR: M=0.00056, SEM=0.00021; REL: M=0.00064, SEM=0.00029; REM: M=0.00063, SEM=0.00023; MDD: M=0.00055, SEM=0.00019).

MVC did not correlate with overall probability to accept ($r=0.005$, $p=0.95$). Adding MVC as an additional covariate (mean corrected) did not change the results for the model-agnostic (main effect of group: $F(3,174)=3.27$, $p=0.023$) or computational results (bias parameter - main effect of group: $F(3,174)=3.16$, $p=0.026$; REM vs REL: $p=0.006$; REL vs HC: $p=0.034$; MDD vs REL: $p=0.047$).

##### Fatigue analysis

To assess whether group differences in acceptance rates could be influenced by fatigue, we conducted two sensitivity analyses. The first analysis focused on the immediate impact of exerting effort on subsequent choice behaviour. For this we included a factor examining the effort exerted on the previous trial, categorising it into conditions of no exertion, low effort exertion (effort levels 0.2 and 0.4) and high effort exertion (effort levels 0.6 and 0.8) on the previous trial. A repeated-measures ANOVA was performed, including effort, reward, and prior-trial-exertion level as within-subject factors, and group as a between-subject factor, with age included as a covariate. Reward and effort levels were categorized as "low" (levels 3 and 6 in reward, and 0.2 and 0.4 in effort) and "high" (levels 9 and 12 in reward, and 0.6 and 0.8 in effort), as the prior-trial-exertion factor lacked sufficient trials to analyse each specific level separately. Out of the original 180 participants, 15 (5 MDD, 3 REM, 1 REL, 6 CTR) had to be excluded due to having no data in at least one cell in this analysis.

Similar to the primary analysis reported in the main text, there were significant main effects of reward (F(1, 160)=362.82, p<0.001) and effort (F(1, 160)=424.27, p<0.001), and a significant effort-by-reward interaction (F(1, 160)=275.33, p<0.001). There was no main effect of previous effort exertion or any interaction with this factor (all ps>0.05; Figure S6A). Consistent with the primary analysis, there was a significant main effect of group on acceptance rates (F(1, 160)=2.80, p=0.04; REM<REL, p=0.009; REM<CTR, p=0.035).

For the second sensitivity analysis, we investigated how cumulative fatigue might influence choice behaviour, over the course of the task. To achieve this, we divided the task into three blocks (the minimum feasible number, given the number of trials) and conducted a repeated-measures ANOVA including reward level (low, high), effort level (low, high), and block (1, 2, 3). Group was a between-subject factor, with age included as a covariate.

Similar to the primary analysis reported in the main text, there were significant main effects of reward (F(1, 175)=416.47, p<0.001) and effort (F(1, 175)=431.53, p<0.001), and a significant reward-by-effort interaction (F(1, 175)=292.96, p<0.001). There was also a significant main effect of block (F(1.91, 333.36)=4.75, p=0.01) and a significant block-by-effort interaction (F(2,350)=5.74, p=0.004). This interaction was driven by lower acceptance rates in the third versus both the first (p=0.002) and second (p=0.011) blocks for high-effort trials (Figure S6B). Thus, cumulative fatigue affected choices particularly when the required effort was high. Importantly, the main effect of group remained significant (F(1, 175)=3.24, p=0.023; MDD<REL, p=0.013; REM<REL, p=0.007) and there were no significant interactions between group and any of the factors (all p>0.05), including block (F(5.72, 333.36)=0.87, p=0.52).

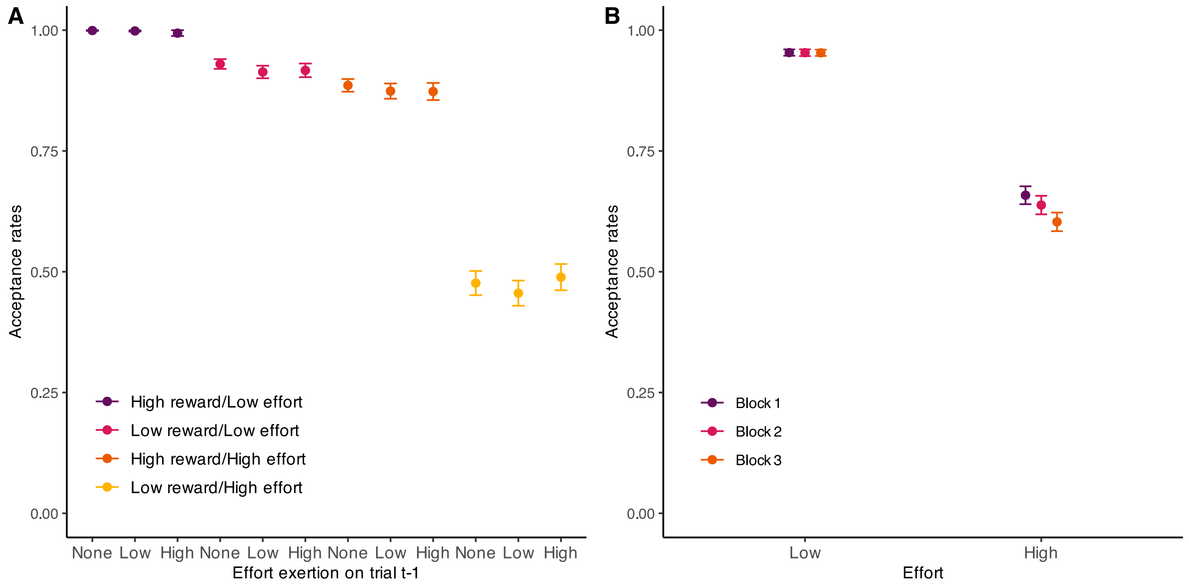


***Figure S6: Fatigue effects.*** *(A) The effect of effort exertion on the previous trial and (B) block when effort levels are low and high on acceptance rates. Figure is showing raw data but analyses were done on transformed data. Error bars represent standard error of the mean.*

#### Computational results: Model comparison

##### *Pilot study*

In the Pilot study, we observed that a specific family of models (Figure S7) was able to account for participants' decisions and performed well in the model comparison procedure. This family always included an intercept parameter (K), which accounts for general tendency to accept offers and a linear reward sensitivity parameter (LinR). The difference between the winning models was in the implementation of the effort sensitivity profile. The best performing models incorporated effort as either a single linear parameter (LinE), a single quadratic effort parameter ($E^2$; similar to[12]), or both linear and quadratic effort parameters (LinE + $E^2$). A variant within this family included an additional noise (or lapse: L; eq. 5) parameter, to account for random responding. The higher the noise term, the more likely a participant will respond at random on a given trial.

$$Eq. 5\text{: Observation} = \left(\theta_{noise} \times \frac{1}{2}\right) + [(1-\theta_{noise}) \times \ Bernoulli(eq.4)]$$

There was positive evidence (difference in k-fold CV score=5.42) for the winning model (LinR+LinE+$E^2$+K), which included the quadratic effort parameter, compared to the second-best model (LinR+LinE+K+L). A similar model that included the noise term (winning model +L), had slightly better evidence than the winning model. However, this was driven by a single participant who was estimated to respond at random on ~30% of trials. We therefore opted to use the simpler model without the noise parameter, as this aided comparison between the two studies (individual estimated parameters were almost identical between these models, all rs>0.99).

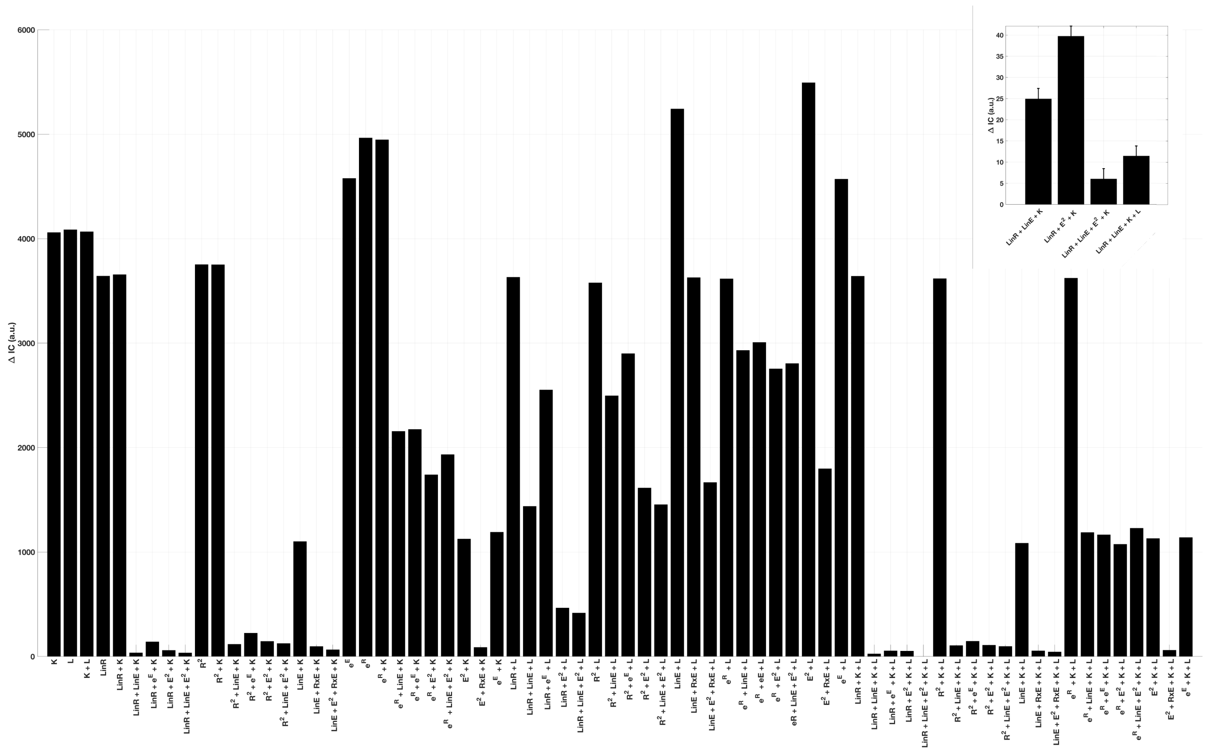

***Figure S7: Model comparison using K-fold cross-validation for the Pilot study.*** *Seventy models were fitted, tested and compared against each other. Inset figure displays a family of models that consistently displayed good fit with low overfitting (lowest bars): Linear reward sensitivity (LinR) + Linear effort sensitivity (LinE) + intercept (K) + with/out Quadratic effort sensitivity (QuadE) + with/out Noise parameter (L).*

*Case-control study*

In the Case-control study, we found that a similar family of models (Figure S8) was present within the top performing models to that observed in the Pilot study. These models again included an intercept (K), linear reward sensitivity (LinR), and various implementations of effort sensitivity profiles (LinE, $E^2$, or both). However, models including the lapse parameter performed poorly, as no participant displayed high levels of random responding.

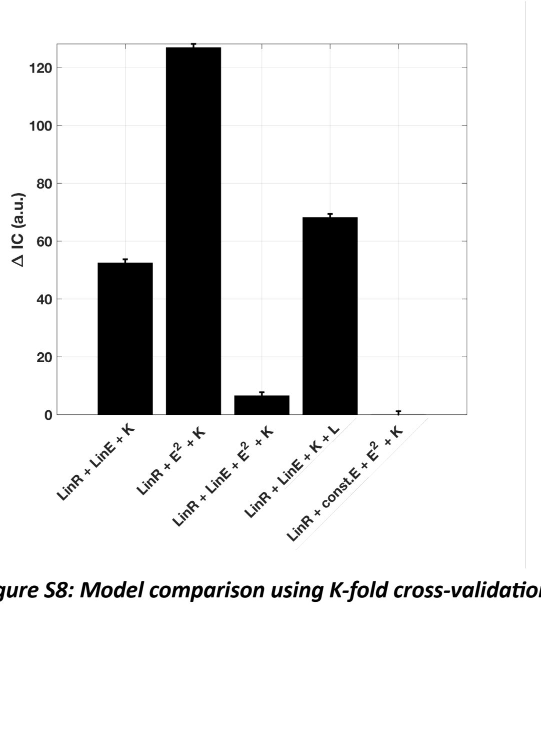


***Figure S8: Model comparison using K-fold cross-validation for the top performing models in the Case-control study.***

During model checking for both studies, we identified that the LinE and $E^2$ parameters were substantially positively correlated (r=0.8), suggesting a trade-off which could lead to poor parameter identifiability due to shrinkage. Moreover, especially in the Case-control study, the range of the LinE term was extremely low [-13.9 to -14.6], suggesting that variability in this parameter did not contribute substantially to decisions. Therefore, we tested an additional model in which the LinE term was constrained to that of the average LinE parameter estimated from the Pilot study (ConstE=-15). This allowed us to avoid parameter trade-off by simplifying the model, and improve parameter identifiability. This value of ConstE also happens to scale the effort levels (20%/40%/60%/80%) to the same range as the reward levels (scaled effort values: -3, -6, -9, -12). There was strong evidence (difference in k-fold CV score=6.52) for the model including the fixed ConstE term (LinR+ConstE+$E^2$+K) compared to the second-best model (LinR+LinE+$E^2$+K). Therefore we implemented this model in the Case-control study analysis.

#### References


1. Sheehan DV, Lecrubier Y, Harnett Sheehan K, et al. The validity of the Mini International Neuropsychiatric Interview (MINI) according to the SCID-P and its reliability. *European Psychiatry*. 1997/01/01/ 1997;12(5):232-241. doi:https://doi.org/10.1016/S0924-9338(97)83297-X
2. Gershon ES, DeLisi LE, Hamovit J, et al. A controlled family study of chronic psychoses: schizophrenia and schizoaffective disorder. *Archives of general psychiatry*. 1988;45(4):328-336.
3. Maxwell ME. Family Interview for Genetic Studies (FIGS): a manual for FIGS. *Bethesda, MD: Clinical Neurogenetics Branch, Intramural Research Program, National Institute of Mental Health*. 1992;
4. Hamilton M. Development of a rating scale for primary depressive illness. *Br J Soc Clin Psychol*. Dec 1967;6(4):278-96. doi:10.1111/j.2044-8260.1967.tb00530.x
5. Vehtari A, Gelman A, Gabry J. Practical Bayesian model evaluation using leave-one-out cross-validation and WAIC. *Statistics and Computing*. 2017/09/01 2017;27(5):1413-1432. doi:10.1007/s11222-016-9696-4
6. Watanabe S. Asymptotic equivalence of Bayes cross validation and widely applicable information criterion in singular learning theory. *Journal of machine learning research*. 2010;11(12)
7. Kass RE, Raftery AE. Bayes Factors. *Journal of the American Statistical Association*. 1995/06/01 1995;90(430):773-795. doi:10.1080/01621459.1995.10476572
8. Gelman A, Hwang J, Vehtari A. Understanding predictive information criteria for Bayesian models. *Statistics and Computing*. 2014/11/01 2014;24(6):997-1016. doi:10.1007/s11222-013-9416-2
9. Gelman A, Rubin DB. Inference from Iterative Simulation Using Multiple Sequences. *Statistical Science*. 11/1 1992;7(4):457-472. doi:10.1214/ss/1177011136
10. Gabry J, Simpson D, Vehtari A, Betancourt M, Gelman A. Visualization in Bayesian Workflow. *Journal of the Royal Statistical Society Series A: Statistics in Society*. 2019;182(2):389-402. doi:10.1111/rssa.12378
11. Gelman A, Hill J. *Data Analysis Using Regression and Multilevel/Hierarchical Models*. Analytical Methods for Social Research. Cambridge University Press; 2006.
12. Chong TT, Apps M, Giehl K, Sillence A, Grima LL, Husain M. Neurocomputational mechanisms underlying subjective valuation of effort costs. *PLoS Biol*. Feb 2017;15(2):e1002598. doi:10.1371/journal.pbio.1002598
